# AI-assisted Discovery of an Ethnicity-influenced Driver of Cell Transformation in Esophageal and Gastroesophageal Junction Adenocarcinomas

**DOI:** 10.1101/2022.01.30.478408

**Authors:** Pradipta Ghosh, Vinicius J. Campos, Daniella T. Vo, Caitlin Guccione, Vanae Goheen-Holland, Courtney Tindle, Guilherme S. Mazzini, Yudou He, Ludmil Alexandrov, Scott M. Lippman, Richard R. Gurski, Soumita Das, Rena Yadlapati, Kit Curtius, Debashis Sahoo

## Abstract

Although Barrett’s metaplasia of the esophagus (BE) is the only known precursor lesion to esophageal adenocarcinomas (EACs), drivers of the metaplasia→dysplasia→neoplasia cascade in the esophagus remains incompletely understood. Using an AI-guided network transcriptomics approach, in which EAC initiation and progression is modeled as networks to simplify complex multi-cellular processes, we first predict cellular continuum states and disease driving processes with an unprecedented degree of precision. Key AI-guided predictions are subsequently validated in a human organoid model and patient-derived biopsies of BE, a case-control study of genomics of BE progression, and in a cross-sectional study of 113 patients with BE and EACs. We find that all EACs must originate from BE, pinpoint a CXCL8/IL8↔neutrophil immune microenvironment as a driver of cellular transformation in both EACs and gastroesophageal junction-ACs. This driver is prominent in Caucasians (Cau), but notably absent in African Americans (AAs). Network-derived gene signatures, independent signatures of neutrophil processes, CXCL8/IL8, and an absolute neutrophil count (ANC) are associated with risk of progression. SNPs associated with ethnic changes in ANC modify that risk. Thus, findings define a racially influenced immunological basis for cell transformation and suggest that benign ethnic neutropenia in AAs may serve as a deterrent to BE→EAC progression.

**BRIEF SUMMARY:** Esophageal adenocarcinoma (EAC) is a highly lethal cancer among Caucasians, while African Americans are somewhat protected; what factors drive transformation with racial disparity remain unknown. AI-enabled creation of the first computational map of neoplastic progression in the esophagus built and validated using transcriptomic datasets from diverse cohorts of human samples pinpointed CXCL8↔neutrophil tumor immune-microenvironment as a racially influenced driver of EACs and GEJ-ACs. Computational tools pinpoint a racially influenced driver of cell transformation during BE→EAC progression; in doing so, it reveals new novel biology, informs disease modeling, therapeutic strategies, and biomarkers.

**LAY SUMMARY:** By modeling diseases as networks, this work unravels a fundamental race-influenced immunologic driver of cell transformation in adenocarcinomas of the esophagus and the gastroesophageal junction.

## INTRODUCTION

EACs are devastating cancers with high mortality (5% 5-y survival^1, 2^). BE is the only known precursor lesion; it is not fully understood why only a small proportion of BE lesions progress to EACs^3^. Also unknown is why EACs display racial disparity^4–6^; African Americans (AAs) are ~4-5-fold less likely to get it than Caucasians (Cau). In fact, risk factors for EAC, e.g., long-segment BE and dysplastic-BE are also less frequent in AA than in Cau^7^.

As to what propels the progression of BE to EACs, approaches to interrogate the genome (whole genome/exome sequencing, etc.) have revealed that BE displays early genomic instability, i.e., in copy number variation that is detectable later in EACs and have revealed patterns of progression (either ‘born bad’, gradual or catastrophic^3^) that may help to describe the evolution of EACs. Despite the insights gained, these genomic insights are yet to translate into prognostic biomarkers of the risk of BE→EAC progression, and answers to fundamental questions, i.e., what drives cellular transformation in BE lesions and if some of those drivers are racially influenced have remained elusive.

Modeling human diseases as networks simplify complex multi-cellular processes, helps identify patterns in noisy data that humans cannot find, and thereby improves precision in prediction. Although transcriptomic network studies in most cancers is a mainstream approach, the field of BE/EAC is notable for its absence. Here we present a network-based approach that uses artificial intelligence (AI) to identify continuum states (of tissues, cell types and processes, and signaling events and pathways) during the process of disease initiation and progression. Once built, the network model then guides subsequent validation (**Figure 1**). We demonstrate how this approach aid in the discovery of fundamental progressive timeseries events underlying complex human diseases and exploit such insights to deliver important insights. More specifically, we use this approach to not only identify key drivers of the metaplasia→dysplasia→neoplasia cascade in the esophagus, but also validate them in a recently described organoid model of BE^8^, tissues and in a cross-sectional retrospective study on a cohort of patients with BE and EAC and analyze if/how race and/or ethnicity may impact these drivers (see study design in **Figure 1**).

**Figure 1.**
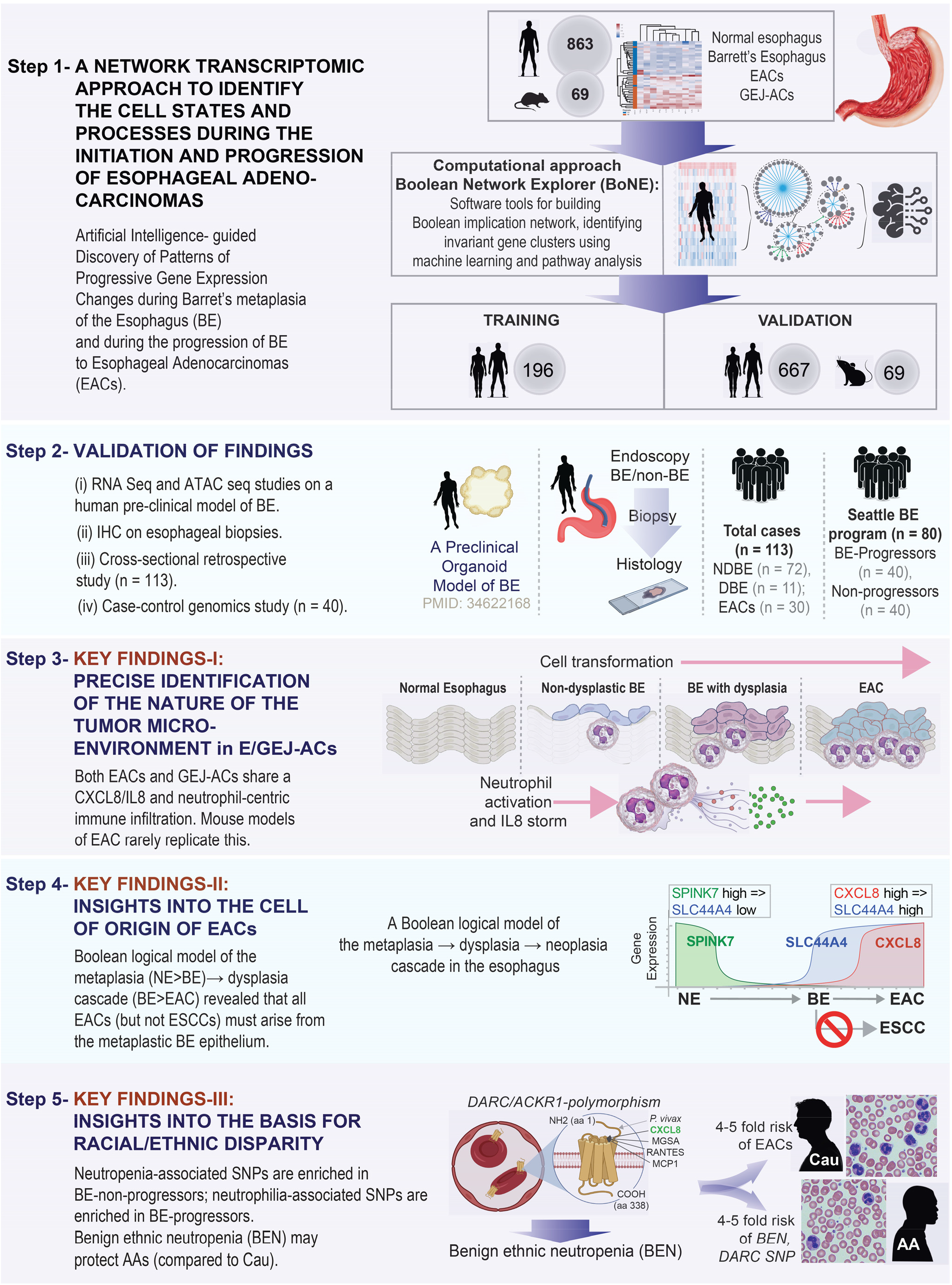
Study Design. *Step 1:* A database containing 932 gene-expression data from both human (n = 863) and mouse (n = 69) samples was mined to build a validated Boolean implication network-based computational model of disease continuum states during the metaplastic→dysplastic→neoplastic cascade in the squamous epithelial lining of the esophagus. Paths, clusters, and list of genes in the network-based model are first prioritized by machine learning approaches and used subsequently to discover cell types and cellular states that fuel the cascade. *Step 2:* Schematic displays the four different models and approaches were used to validate the network-derived findings. *Step 3-5:* The network-based model reveals three key findings.

## RESULTS AND DISCUSSION

### An AI-assisted study design

To identify cellular continuum states during pre-cancer to cancer progression, we chose a Boolean approach to building transcriptomic networks^9^. This approach has been used to create maps of evolving cellular states along any disease continuum and identify hitherto unforeseen cellular states in diverse tissues and contexts with high degrees of precision. For example, it helped pinpoint branchpoints in B/T cell differentiation^10,11^, define progenitor cell hierarchy in blood^12–16^, normal and neoplastic cell states in colorectal cancers^17,18^, bladder cancers^19–22^, and prostate cancers^23,24^, and identify NK cell exhaustive states^25^, universal cell proliferative^26^ and macrophage^27^ markers, and cell states in the mucosal barrier in IBD^28^. The superiority of the Boolean approach over other conventional approaches in accurately modeling gene expression changes and predicting clinical outcomes has been shown before^17,18,20, 23^ in head-to-head comparisons (see *Methods*). Because the Boolean approach relies on invariant relationships that are conserved despite heterogeneity in the samples used for the analysis, which often represent maximum possible diversity, i.e., the relationships can be thought of as general relationships among pairs of genes across all samples irrespective of their origin (normal or disease), laboratories/cohorts, different perturbations, and sometimes in multiple species including human, mouse and rat, and hence, considered conserved invariants. It is assumed that such ‘invariants’ are likely to be fundamentally important for any given process.

We used the Boolean approach to build maps of continuum states first during metaplastic progression in the normal esophagus (NE→BE), and subsequently during neoplastic transformation of the metaplastic epithelium (BE→EAC). Gene signatures were identified from each map, independently, using machine learning approaches. Signatures were validated in numerous independent cohorts (see **Figure 1**; top), prior to their use in validation studies; the latter included various experimental approaches on human tissues or tissue-derived organoids (**Figure 1**; middle). Map-derived gene signatures in conjunction with multiple independent gene signatures were used as precise and objective tools to navigate new biology, and to formulate and rigorously test new hypotheses, which led to a few notable findings (**Figure 1**; bottom).

### A Boolean map of metaplastic progression in the esophagus

We used Boolean Network Explorer (*BoNE*)^29^ to first create a model of progressive gene regulatory events that occur during metaplastic transition (**Figure 2A**). For model training and development, we used the largest (to our knowledge) well annotated transcriptomic dataset [n = 76: <u>GSE100843</u>^30^] derived from BE and proximal matched normal mucosa from squamous esophagus from 18 BE patients. Gene expression patterns were first simplified into ‘clusters’ within which each gene is equivalent to one another (**Figure 2B**). The clusters (nodes) were connected to one another (starting with the largest) based on the pattern of relationships between the clusters (edges), conforming to one of the six possible Boolean implication relationships (BIRs; **Figure 2C**). These efforts helped chart numerous Boolean paths (**Figure 2D**, *left*) within a network with directed edges (**Figure 2E**). Each cluster was then evaluated for whether they belong to the healthy esophagus or diseased side (BE) depending on whether the average gene expression value of a cluster in heathy samples is up or down, respectively. Each path of connected gene clusters indicates a certain hierarchy in gene expression events, which translates to a progressive series of gene down/upregulation events, predicted to occur in sequence during the metaplastic process (**Figure 2D,** *right*).

**Figure 2.**
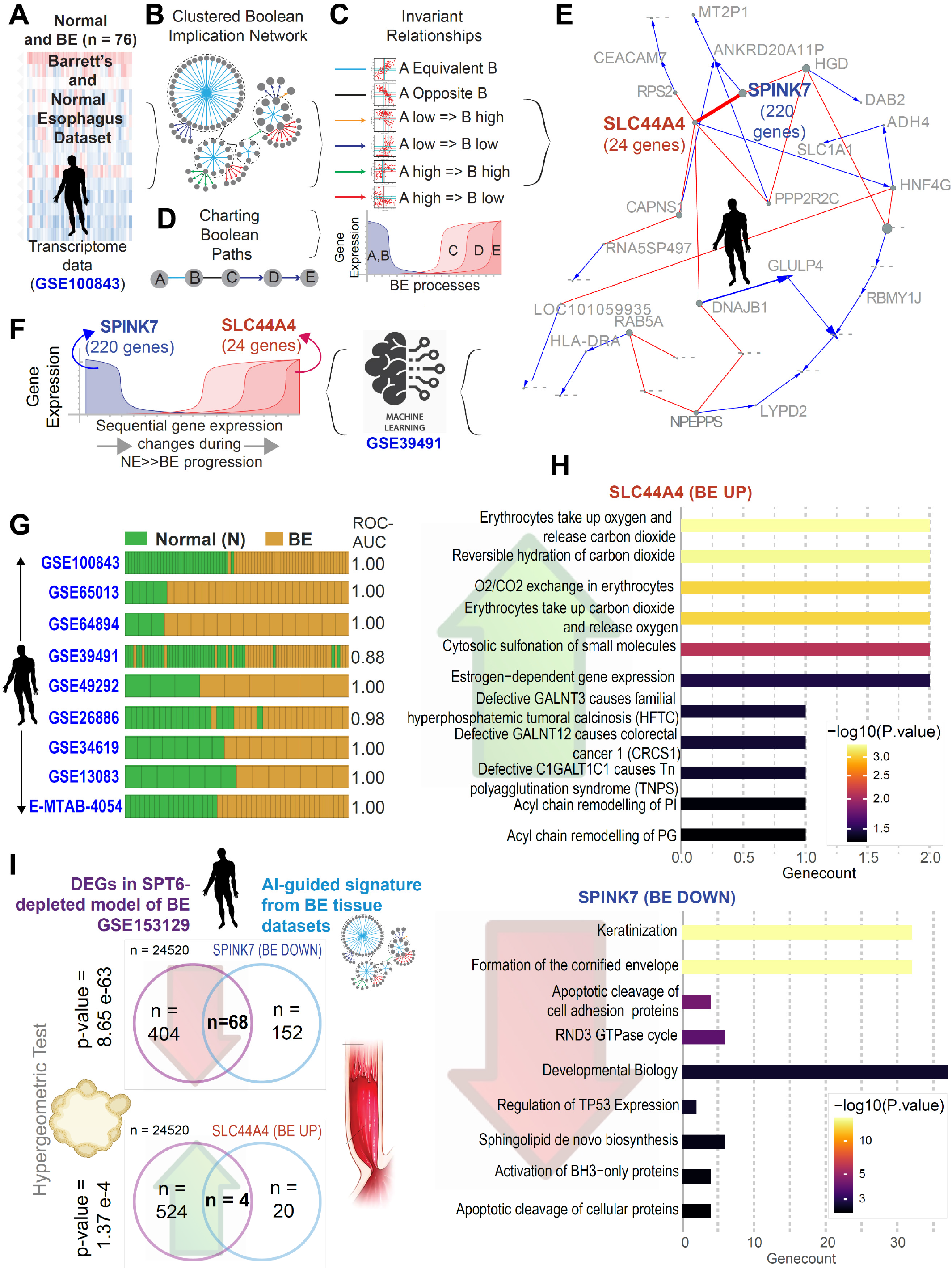
Generation and validation of Boolean network map of Barrett’s metaplasia of the esophagus. **A-D**. Schematics outline the workflow and training datasets used to create a Boolean map of changing expression of gene clusters during normal esophagus (NE) to BE transition using *BoNE*^29^. We applied *BoNE* to analyze dataset GSE100843 (A). The Boolean network (B) contains the six possible Boolean relationships (C) in the form of a directed graph with genes having similar expression profiles organized into clusters, and relationships between gene clusters represented as color-coded edges (B). Boolean cluster relationships are used to chart disease paths (D-left); and individual gene expression changes along a boolean path, illustrating gene expression dynamics within the normal to BE to EAC continuum (D-right). **E**. Graph showing invariant patterns (down or up-regulation) of gene expression changes during NE>BE progression. Two clusters of genes (*SPINK7* and *SLC44A4*) identified by machine learning are indicated in bold. **F**. Gene clusters in E were filtered through a second dataset [GSE39491] to refine the gene signature (left). The resultant signature involves progressive downregulation of *SPINK7*-cluster with a concomitant upregulation of *SLC44A4*-cluster (*right*). **G**. Bar plots showing sample classification accuracy across diverse BE validation datasets, with corresponding ROC-AUC values. **H**. Reactome pathway analysis of *SPINK7-* and *SLC44A4*-clusters were performed to identify the signaling pathways and cellular processes that are enriched during metaplastic progression in esophagus. **I**. Gene signatures UP- or DOWN-regulated in BE-map (which is built using tissue transcriptomics) are compared against differentially expressed genes (DEGs) in our recently published SPT6-depleted organoid model of BE^8^. Hypergeometric statistical analyses show significant overlaps in both UP and DOWN-regulated genes.

We next introduced in *BoNE* machine learning that seeks to identify which of the gene clusters (nodes) connected by Boolean implication relationships (edges) are most optimal in distinguishing healthy from diseased samples. *BoNE* computes a score that naturally orders the samples; this score can be thought of as a continuum of states. Among all possible permutations and combinations, a set of two clusters emerged as most robust, which was further refined by an additional filtering step through a second ‘training dataset’ [<u>GSE39491</u>^31^; see **<u>Supplemental Information 1</u>**] which is comprised of BE and normal esophageal matched samples from 43 patients. Both training datasets were analyzed independently throughout the process. The resultant model of metaplastic transition pinpointed a time series of metaplasia (BE)-associated invariant events, in which downregulation of expression of 220 genes (*SPINK7*-cluster; **Figure 2F**) was invariably associated with a concomitant upregulation of 24 genes (*SLC44A4*-cluster, **Figure 2F**) in all samples in the training datasets. *SPINK7* (serine peptidase inhibitor, kazal type 7), is a key checkpoint in the esophageal keratinocyte stem cell, which regulates mucosal differentiation, barrier function, and inflammatory responses^32^. *SLC44A4* encodes a specific high-affinity regulated carrier-mediated uptake system for TPP in human colonocytes, involved in regulation of microbiota-generated thiamine^33^. The pattern of gene expression signature was sufficient to classify samples in seven independent validation cohorts and performed consistently well when doing so (ROC AUC 0.88-1.00; **Figure 2G**). For a complete list of genes in these clusters see **<u>Supplemental Information 2</u>**.

Reactome pathway analysis of the upregulated *SLC44A4-* and downregulated *SPINK7*-clusters along the path continuum revealed the most important biological processes that they control (**Figure 2H**). The downregulated pathways (**Figure 2H;** *bottom*) were cellular processes that are inherently associated with squamous epithelium, e.g., keratinization, cornified envelope formation, as expected. Other notable changes were TP53 expression and cell-cell adhesion proteins. These findings are consistent with emerging evidence from numerous independent studies which agree that aberrant TP53 IHC highly correlated with *TP53* mutation status (90.6% agreement) and was strongly associated with higher risk of neoplastic progression regardless of the presence/absence of dysplasia^34–36^. The findings are also in keeping with the reduction observed by IHC in cell adhesion proteins in BE lesions [E-cadherin, P-cadherin and the catenins which serve as adaptor proteins that enable the cadherins to achieve cell adhesion^37^]. The most notable cellular processes that were upregulated (**Figure 2H;** *top*) were related to oxygen delivery to the tissue, consistent with reports of Warburg and Crabtree effects in BE tissues^38^.

### BE map-derived signatures support an epigenetic cascade with loss of keratinocyte identity

We began by confirming that the network-derived signatures were recapitulated by our recently published^8^ organoid model of BE, i.e., overlaps between upregulated DEGs and the *SLC44A4*-cluster and between downregulated DEGs and the *SPINK7*-cluster were significant (p-value 1.37 e-4 and 8.65 e-63, respectively; **Figure 2I**). This model is unique in that it does not represent established BE; instead, it recapitulates the process of metaplastic transformation from stratified squamous to intestinal epithelium. This model emerged serendipitously during studies interrogating fate determinants of human keratinocyte stem cells using an unbiased siRNA screen approach^39^ (**Figure 3A**); loss of transcription elongation factor SPT6 emerged as a *bona fide* trigger for epithelial transcommitment from stratified squamous epithelium to a ‘intestine-like’ lineage (**Figure 3B**). Mechanistically, depletion of SPT6 (in conjunction with rescue studies) confirmed that such transdifferentiation was primarily due to stalled transcription and downregulated expression of TP63, the master regulator of keratinocyte fate and differentiation^40–43^. Computational approaches confirmed that this phenomenon of *transcommitment* faithfully recapitulates the metaplasia-specific signatures of BE^8^, i.e., differentially expressed genes (DEGs) from nine independent BE datasets were captured in this model (**Figure 3C**). Transient acid exposure downregulated SPT6 and TP63, both transcript and protein^8^. The proposed working model was that acid exposure downregulates SPT6, which in turn inhibits TP63, which is sufficient for a cell fate switch from epidermis to a metaplastic columnar epithelium; the latter is indistinguishable from BE, both morphologically and by gene expression^8^.

**Figure 3.**
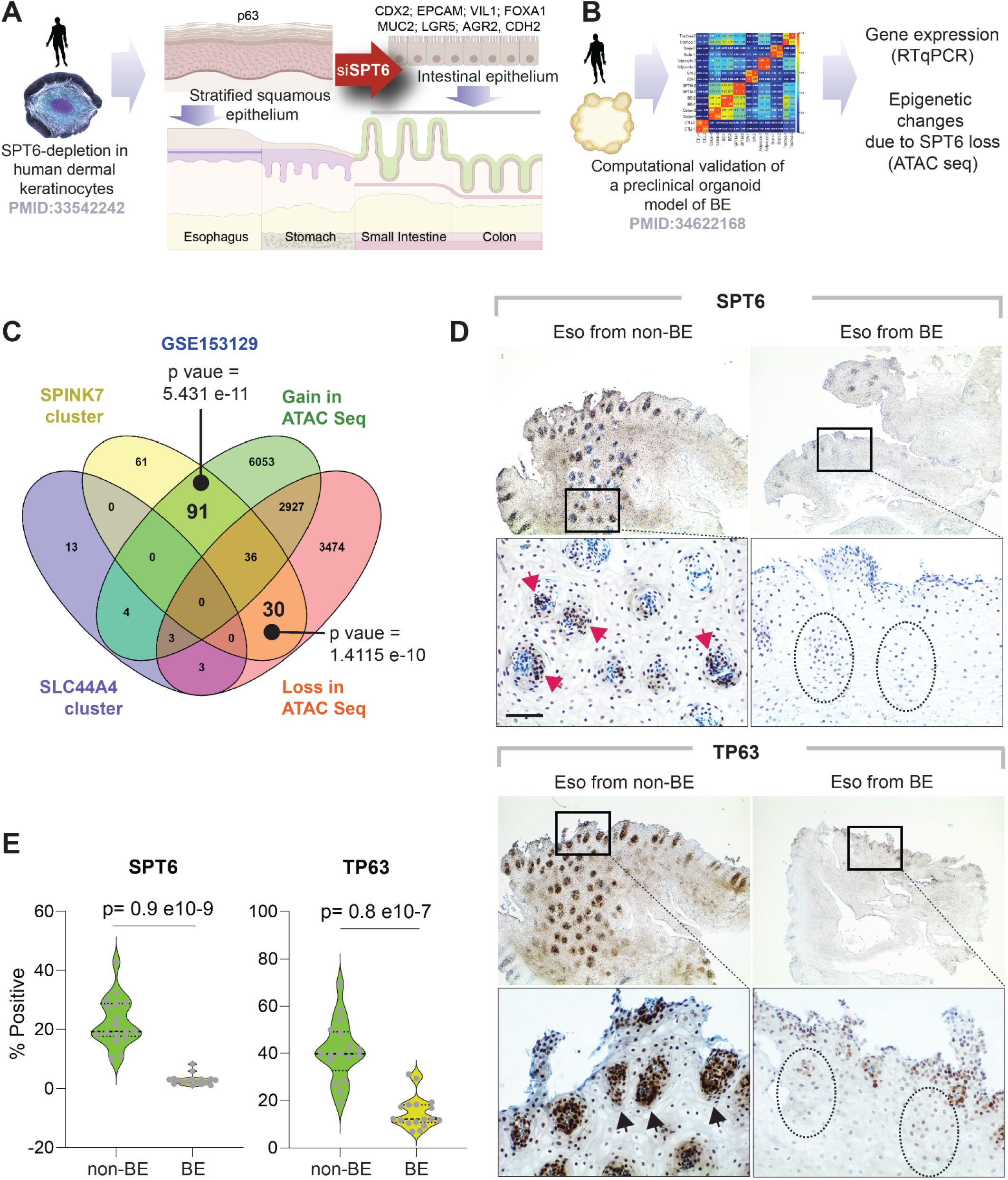
Validation of the Boolean map of Barrett’s esophagus using a human organoid model of squamous metaplasia. **A-B**. Schematic summarizing the key findings in gene expression and epithelial morphology observed and reported earlier ^39^, upon depletion of SPT6 in keratinocyte stem cells by siRNA ^39^. While control keratinocytes formed stratified squamous epithelium, siRNA mediated transient depletion of SPT6 in keratinocytes (SPT6i) grew as ‘intestine-like’ monolayers. RNA seq studies on those monolayers confirmed that this model recapitulates metaplastic gene signatures in BE, and not normal gut differentiation that is observed in the healthy gut lining. **C**. Gene clusters from the BE-map were analyzed for overlap with those affected in the organoid BE model by loss or gain of access due to chromatin remodeling upon SPT6 loss, as identified by ATAC Seq. Only significant *p values,* as determined using hypergeometric analyses, are displayed. **D**. Esophageal biopsies obtained 2 cm above the GEJ from Caucasian males who had BE (Eso from BE) or had not (Eso from non-BE) progressed to BE were analyzed for SPT6 and TP63 expression by IHC. Red and black arrowheads point to crypts staining positive. Interrupted circles highlight crypts with little/no expression. Fields representative from n = 3 subjects are shown; boxed regions above are magnified below; Scale bar = 100 μm. **E**. Violin plots display the % cells positive for staining in regions of interest (ROIs, marked with arrows on left and interrupted ovals on the right), as determined by the ImageJ plug-in, IHC profiler.

We hypothesized that SPT6 deletion may have altered the genome-wide chromatin accessibility to genes (beyond those genes that are targets of *TP63*) in the BE-map derived *SPINK7* and *SLC44A4* clusters. ATAC-seq (Assay for Transposase-Accessible Chromatin using sequencing) studies on the SPT6-depleted BE organoid model confirmed that it is the DOWN-regulated genes in the *SPINK7*-cluster, which includes *TP63*, are significantly impacted when the histone chaperone SPT6 fails to interact with histone H3 and elongate Pol-II to regulate chromatin dynamics and mRNA biogenesis. Findings support our prior conclusions^39^, in that the *SPT6→TP63* axis maintains keratinocyte identity (pathways enriched in the *SPINK7* cluster; **Figure 2H**), and that its loss permits transcommitment to a metaplastic intestine-like fate.

To analyze the translational relevance of these findings, we next asked if the *SPT6→TP63* axis is downregulated in the squamous esophageal lining in patients with BE. We prospectively enrolled patients with or without BE and collected biopsies from the distal esophagus, 2 cm above the GEJ or the BE segment. Immunohistochemistry studies on FFPE-embedded biopsies confirmed that, compared to non-BE subjects, both SPT6 and TP63 proteins were significantly suppressed in the esophageal squamous lining from patients who have progressed to BE (p values 0.8 e-9 and 0.9 e-7, respectively; **Figure 3D-E**). These findings support keratinocyte transdifferentiation as a trigger for BE, as proposed by multiple independent groups^44–47^, but cannot rule out gastric cardia as alternative sites of initiation.

### A Boolean map of EAC identifies an ‘immune paradox’ during cell transformation in EACs and GEJ-ACs

Next, we created a Boolean map of early cellular states that are encountered during BE→EAC transformation (**Figure 4A**). A sequence of gene expression changes emerged: a cluster of 471 genes (*LNX1*-cluster) were downregulated, with a concomitant and sequential upregulation of two clusters totaling another 61 genes (*IL10RA-* and *LILRB3*-clusters) invariably encountered in all samples (**Figure 4A**-*right*; Figure 4B). Machine learning approaches pinpointed the *LILRB3-* and *IL10RA*-clusters as sufficient to classify EACs from BE and do so reproducibly in 4 independent validation cohorts (**Figure 4C**). Because one of the strengths of the Boolean approach is its ability to extract using the logic of asymmetric invariant relations the fundamental timeseries of events in cross-sectional datasets^10^, we saw that the UP-signatures are induced in a sequential manner (**Figure 4B**): While the *IL10RA*-cluster is up in non-dysplastic BE (**Figure 4D**-*left*), LILRB3-cluster is induced predominantly in EACs (**Figure 4D**-*middle*); the composite score of the combined signature shows progressive increase throughout BE→EAC transformation (**Figure 4D**-*right*). The genes in these clusters are listed in **<u>Supplemental Information 3</u>**

**Figure 4.**
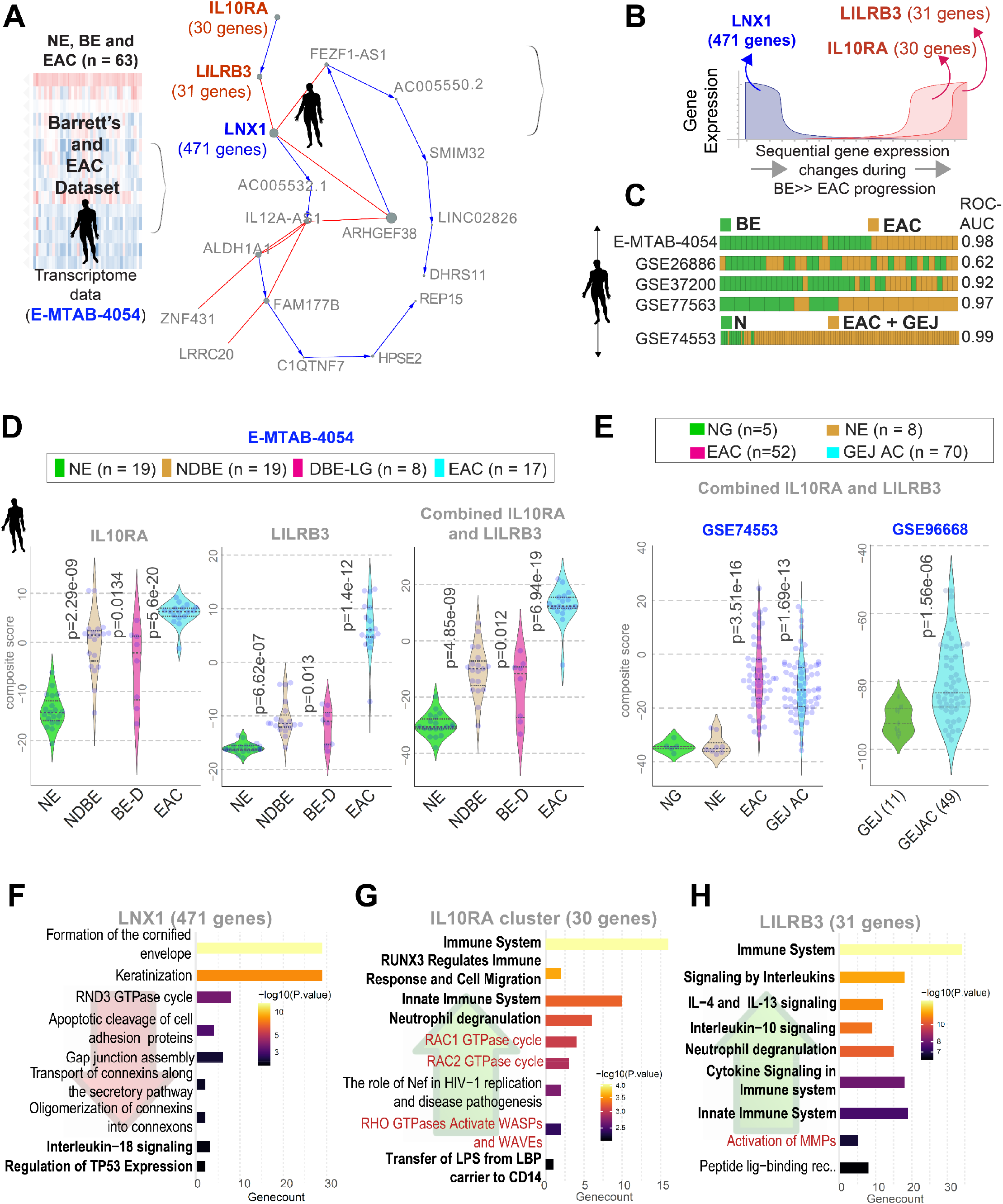
Generation and validation of Boolean network map of BE to EAC progression. **A**. Schematic outlining the workflow and training datasets used to create a network map of gene clusters using our published computational platform, *BoNE*^29^. **B**. Graph showing invariant patterns (down or up-regulation) of gene expression changes during BE>EAC transformation. Three clusters of genes identified by machine learning are indicated. **C**. Bar plots showing sample classification accuracy across diverse validation datasets, with corresponding ROC-AUC values. **D**. Violin plots showing the composite scores of upregulated gene clusters in normal esophagus (NE), non-dysplastic BE (NDBE), dysplastic BE (DBE), and EACs. *P* values indicate comparison of each sample type against the normal esophagus (NE), as determined by Welch’s t-test. **E**. Violin plots showing the composite scores of upregulated gene clusters in normal esophagus (NE), normal gastric (NG), normal GEJ (GEJ) and GEJ-ACs. *P* values indicate comparison of each sample type against the normal esophagus (NE; left, GSE74553) or GEJ (right, GSE96668), as determined by Welch’s t-test. **F-H**. Pathway analyses (reactome.org) of gene clusters reveal cellular processes that are UP or DOWN during cell transformation in 3 clusters derived from the EAC map in B. Red fonts indicate likely epithelial processes.

The degree of induction of the EAC-map derived signatures was indistinguishable in EACs and gastroesophageal junction adenocarcinomas (GEJ-ACs; <u>GSE74553</u>; **Figure 4E**-*left*), in that both tumor samples showed induction of the signature compared to normal esophagus (NE) and gastric (NG) samples. We validated this finding in another independent dataset (<u>GSE96668</u>; n = 60; **Figure 4E**-*right*), which was comprised of 11 normal GEJ tissues and 49 GEJ-ACs.

Reactome pathway analyses of these gene signatures revealed the set of cellular types and states that are progressively gained or lost (**Figure 4F-H**). Beyond the early Rac and Rho activation that were shown to be contributors during EAC initiation and progression^48^ (**Figure 4G**; red font), the overwhelming and progressively increasing processes were that of innate reactive immune response and inflammatory cytokine signaling (**Figure 4G**), which was predominantly neutrophil centric and comprised of chemokines and receptors that specifically target the neutrophils (e.g., *CXCL8/IL8, CXCR1*, and *CXCL2*; see *LILRB3*-cluster in **<u>Supplemental Information 3</u>**). This is followed by an increase of an equally prominent alternative (immunotolerant/suppressive) immune response that is IL10, IL4/IL13-centric (**Figure 4H**). Such immunosuppressive states have indeed been reported in the setting of EAC by others^49^, using a plethora of approaches, ranging from increased IL10 levels in the serum^50^, to confirmed expression of IL10 protein/mRNA in the tissue^51, 52^; all studies agree that higher immunosuppression carries worse prognosis. The upregulation of both reactive and tolerant immune responses, hence, paradoxical immune response, was associated with a concomitant loss of two known tumor suppressive processes in the esophagus: IL18 signaling^53–55^, which plays a suppressive role uniquely in GI tract cancers^56^, and the TP53 pathway (**Figure 4F**). It is noteworthy that loss of TP53 itself is known to orchestrate a distinct immune cell infiltration^57^ and response.

These findings show that the dynamic immunologic landscapes predicted by the map conform to prior knowledge. They also increase confidence in the map’s ability to formulate new hypotheses and make predictions, which we set out to do next.

### A map of cellular continuum states confirms that all EACs must evolve through BE

We used the concept of Boolean invariant logic to create a model that captures both the early (metaplastic) and late (transformation) steps of cellular state change. First, we found that BE signatures (NE>>BE map; **Figure 2D**) are also induced in the EAC samples across diverse cohorts (**Figure 5A**). In fact, we found that the Boolean Implication *SPINK7* high => *SLC44A7* low, which defines metaplastic transition in the normal epithelium (**Figure 2E**), is an invariant relationship in the most diverse global human <u>GSE119087</u> (n = 25955) dataset (**Figure 5B**), suggesting that this pattern may be fundamentally important. Using *SLC44A4* as a seed gene in a dataset that was comprised of normal esophagus, BE and EAC samples, we found that it shares an invariant relationship with one of the genes in the *LILRB3* cluster, *CXCL8. CXCL8* high => *SLC44A4* high in an invariant Boolean implication relationship in normal esophagus (NE), BE and EAC samples where each sample type is mostly confined to one quadrant (NE, bottom-left; BE, top-left; and EAC, top-right). This model suggests that if EACs must originate from the esophagus, they must do so via the metaplastic BE intermediate; only 3 genes could nearly accurately classify the samples (**Figure 5C**) and show progressive expression changes along the continuum (**Figure 5D**). These findings were validated in a second cohort pooled from multiple independent datasets (**Figure S1A**). By contrast, esophageal squamous cell carcinomas (ESCCs; > 400 samples pooled), did not conform to the Boolean logic-based model of the BE→EC continuum, and as expected, the model confirms that ESCCs do not transition through metaplastic BE states (**Figure S1B**). Datasets that include NE, gastric cardia, BE and EACs/GEJ-ACs could not be pooled (due to incompatible array platforms); this hindered our ability to objectively test if BE→EACs originate in NE or from the gastric cardia; nor could we test if GEJ-ACs originate in BE.

**Figure 5.**
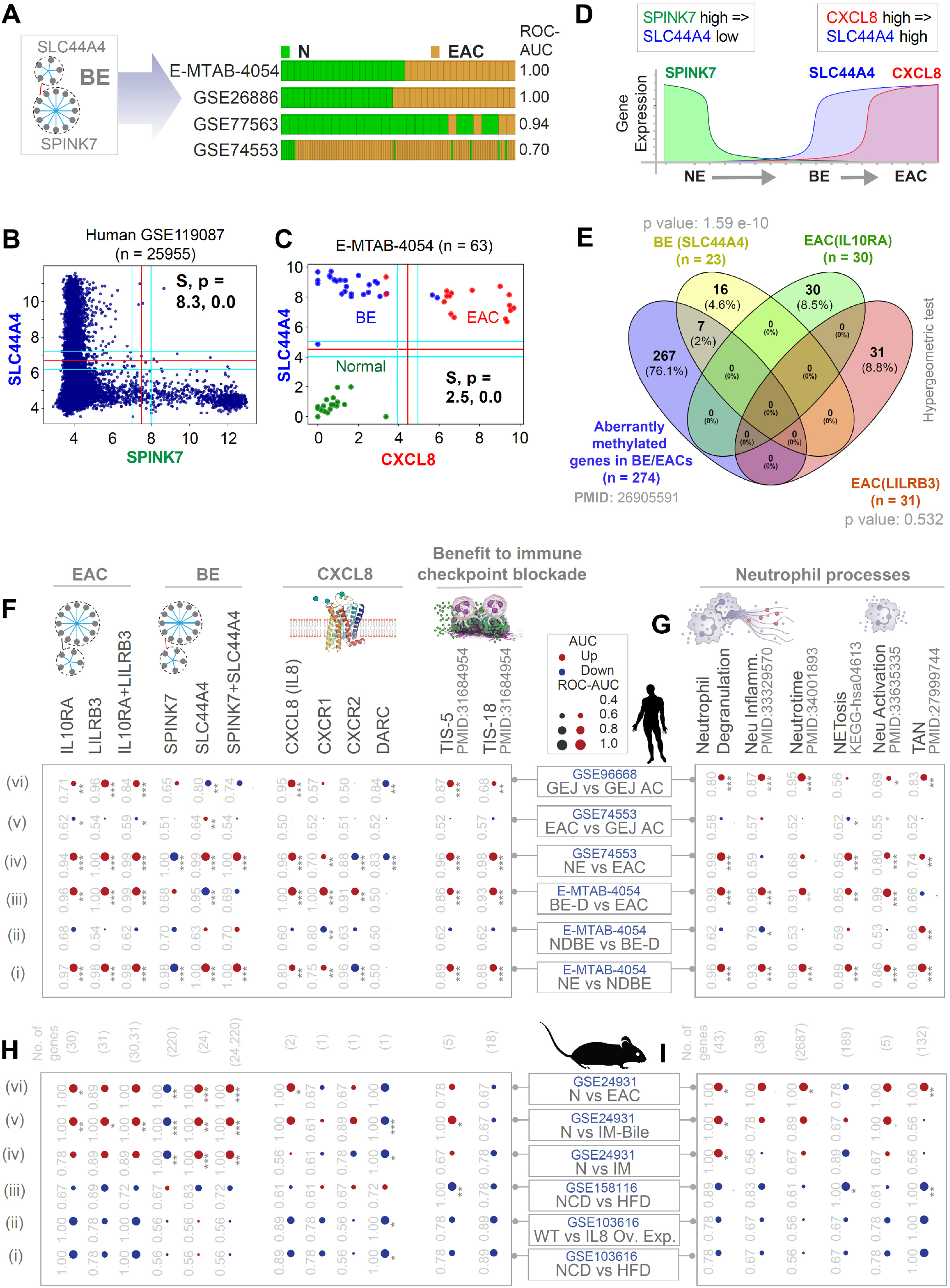
A Boolean logical model of cellular states during NE→BE→EAC progression. **A.** Bar plots showing that BE signatures (NE>>BE map; **Figure 2F-G**) can distinguish normal esophagus (NE) and EAC samples, indicating that these signatures are also induced in EAC samples. Corresponding ROC AUC values are displayed. **B.** A scatterplot for *SPINK7* and *SLC44A4* expression in global human <u>GSE119087</u> (n = 25955) dataset. Boolean Implication SPINK7 high => SLC44A7 low (S=8.3, p=0.0, FDR < 0.001) is an invariant in the most diverse dataset. **C.** A scatterplot for *CXCL8* and *SLC44A4* expression in the <u>E-MTAB-4045</u> dataset. *CXCL8* high => *SLC44A4* high (S = 2.5, p = 0.0, FDR < 0.001) in an invariant Boolean implication relationship in NE, BE and EAC samples where each sample type is mostly confined to one quadrant (NE, bottom-left; BE, top-left; and EAC, top-right). **D.** A schematic to visualize the mathematical model of NE→BE→EAC progression based on MiDReG^10, 11^ analysis using Boolean Implication relationships. Since *CXCL8* high => *SLC44A4* high, and both *CXCL8* and *SLC44A4* are low in NE, the invariant model suggests that BE stage (*SLC44A4* high, *CXCL8* low) must precede *CXCL8* high and *SLC44A4* high. See also **Figure S1** for model validation in an independent cohort of EACs (**S1A**) and for a head-to-head comparison between ESCC vs EAC (**S1B**). **E**. Venn diagram showing the overlaps between the gene clusters from the BE/EAC-maps with the genes reported to be methylated in multiple independent studies (n = 274 / 22178 genes that were tested in total) using Illumina’s Infinium HumanMethylation27 BeadChip microarray. Only significant *p values*, as determined using hypergeometric analyses, are displayed. **F-G**. Human EAC immune microenvironment is visualized as bubble plots of ROC-AUC values (radius of circles are based on the ROC-AUC) demonstrating the direction of gene regulation (Up, red; Down, blue) for the classification of samples (gene signatures in columns; dataset and sample comparison in rows). P values based on Welch’s T-test (of composite score of gene expression values) are provided using standard code (‘‘ p> 0.1, ‘.’ p <= 0.1, * P <= 0.05, ** P <= 0.01, *** P <= 0.001) next to the ROC-AUC. Panel F displays the classification of NE, non-dysplastic BE (NDBE), dysplastic BE (DBE), EAC, GEJ-AC based on signatures derived from the BE>>EAC map (upregulated gene clusters *IL10RA* and *LILRB3*), NE>>BE map (downregulated *SPINK7*-cluster, and upregulated *SLC44A4*-cluster), CXCL8/IL8 and its three receptors (CXCR1/2 and DARC), and tumor inflammation signatures that predict anti-PD1 Rx (TIS-5, TIS-18) in two independent dataset (<u>E-MTAB-4054</u>, <u>GSE74553</u>). Panel G displays the classification of same samples in F based on neutrophil signatures (NETosis, Degranulation, Inflammation, Neutrotime) in <u>E-MTAB-4054</u>, and <u>GSE74553</u>. Violin plots for selected neutrophil signatures across NE, NDBE, DBE, EAC, GEJ-AC samples are displayed in **Figure S2**. See also **Figure S3** for extended data on the relationship between neutrophil and EAC signatures with the TIS signature. **H-I**. Murine BE-EAC immune microenvironment was analyzed using the same set of signatures and visualized as bubble plots exactly as in F-G. IM, intestinal metaplasia. NCD, normal chow diet; HFD, high fat diet; Ov Exp, overexpression. Violin plots for selected signatures are displayed in **Figure S4.**

We analyzed the overlap between our network-derived signatures for BE (**Figure 2E-F; <u>Supplemental Information 2</u>**) or EAC (**Figure 4A; <u>Supplemental Information 3</u>**) and the 274 genes that were reported as aberrantly methylated in 6 independent EAC and/or BE methylation studies (i.e., an integrated methylome of BE; summarized in Krause et. al^58^). These genes can separate NE from EAC/BE, but not EAC from BE, indicating that the same clusters of methylation profiles exist in BE and EAC^58^. Significant overlaps were seen between the 274 genes and the BE-associated *SLC44A4*-cluster (p value 1.59e-10), but not the EAC-associated *LILRB3-* and *IL10RA*-clusters (**Figure 5E**), suggesting that the methylome in EAC is imprinted early during evolution through BE.

### Two ‘waves’ of an IL8↔neutrophil-centric inflammation is encountered during cell transformation

We noted that among the genes that were upregulated during cell transformation (i.e., the BE→EAC map; **Figure 4A**) the *LILRB3*-cluster was enriched for one major cytokine/chemokine pathway and its target immune cell, neutrophils. It contained the *CXCL8* (henceforth, IL8) gene and the receptor for IL8, *CXCR1* (see complete gene list in **<u>Supplemental Information 3</u>**); IL8 was sufficient to classify BE and EAC (**Figure 5C-D**). Discovered as the first chemokine activator of neutrophils, IL8 displays a distinct target specificity for the neutrophil, with only weak effects on other blood cells^59–61^. The *LILRB3*-cluster also contained *CXCL2;* the CXCL2-CXCR2 axis helps in the recruitment of tumor-associated neutrophils^62^.

We asked, how this IL8↔neutrophil centric inflammation varies during the metaplasia-dysplasia-neoplasia cascade by comparing pairwise each sequential step, i.e., normal esophagus (NE) vs non-dysplastic BE (NDBE), NDBE vs dysplastic BE (BE-D), BE-D vs EAC. The induction of BE and EAC map-derived signatures, as well as IL8 and its two signaling receptors (CXCR1/2) (**Figure 5F**) and numerous pathologic neutrophil processes (**Figure 5G; Figure S2**) was significant in EACs (**Figure 5F-G**, *row iv*) and in GEJ-ACs (**Figure 5F-G**, *row vi*). The patterns of induction of all the gene signatures (see **<u>Supplemental Information 4</u>** for gene lists) were virtually indistinguishable between EACs and GEJ-ACs (**Figure 5F-G**, *row v* and **Figure S2C-F-I-L**). Upregulation was observed in two phases: Early during metaplastic transformation from NE to NDBE (*row i*) and later during transformation from BE-D to EACs (*row iii*), but not during BE to BE-D (*row ii*) (see also **Figure S2** for violin plots). These results show that the normal→metaplasia→dysplasia→neoplasia cascade is associated with a staircase waveform of IL8 and neutrophil processes. Because the IL8↔neutrophil axis is known to alter tumor immune landscape^63^ and increased genomic instability^64^, we conclude that a biphasic IL8↔neutrophil interplay is a specific association, and presumably, a potential driver of cell transformation along the BE↔EAC continuum.

### Gene signatures reveal pro-tumor neutrophil infiltrates, benefit from immune checkpoint therapies

We next asked if the increased neutrophil processes were reflective of N1 or N2 profile of neutrophils, which represent immunostimulating (anti-tumor) and immunosuppressive (pro-tumor) subsets, respectively, in cancer^65^. To this end, we analyzed a tumor-associated neutrophil (TAN^66^) signature that measures the protumor N2 TANs. Both EACs (**Figure 5G**-*last column, row iv*) and GEJ-ACs (**Figure 5G**-*last column, row vi*) were associated with an induction of TAN signature. Similarly, upregulation in TAN signature was noted in pairwise comparisons of NE vs NDBE (**Figure 5G**-*last column, row i*) and NDBE vs DBE (**Figure 5G**-*last column, row ii*). ROC AUC and p values are displayed for each pairwise comparison in **Figure 5G**.

The induction of pro-tumorigenic TAN signatures was associated also with the pan-cancer marker of adaptive immune resistance, an 18-gene “tumor inflammation signature” (TIS^67, 68^) and its 5-gene subset (that remained significantly higher in responders after correction for multiple testing^67^) (**Figure 5F***; last two columns*). Higher TIS was demonstrated to retrospectively predict benefit of anti-PD-1 therapy in various cancers^67, 68^. These findings are consistent with reports of higher TIS score in EACs^69^, regardless of treatment status and that Pembrolizumab was found to be effective in esophageal and GEJ cancers (KEYNOTE-590 trial^70,71^ and reviewed in^72^). EAC signatures, neutrophil processes and TIS signatures positively and strongly correlated across all EAC and GEJ-AC datasets analyzed (*r* ranging from 0.8-0.99 or TIS vs EAC signatures; **Figure S3**). These findings suggest that pro-tumorigenic neutrophils may drive adaptive immune resistance.

### Currently available animal models of BE→EAC transformation rarely recapitulate human disease

Animal models of diseases have both merits and limitations^73, 74^. Because ‘mice are not men’^75, 76^, especially when it comes to their innate immune system^76, 77^, and EACs and GEJ-ACs are associated with a prominent immune signature, we asked how well currently available EAC models recapitulate the human disease. To model how EAC-associated risk factors, i.e., obesity/BMI and IL8-induction enhance cell transformation, mice challenged with high-fat diet (HFD^78–80^) or overexpressing IL8^80^ have been developed. Neither model induce our network-derived BE/EAC signatures (**Figure 5H**, *row i-iii*; **Figure S4A-C**); nor did they display induction of the neutrophil signatures we observed in human tissues (compare human-**Figure 5G** with murine-**Figure 5I**, *row i-iii*). The signatures were, however, induced in a transgenic interleukin1-β (IL1β)-overexpression model (<u>GSE24931</u>; **Figure 5H-I**; *rows iv-vi;* **Figure S4D-F**); in this model, the human IL1β cDNA was inserted downstream of an Epstein-Barr virus (ED-L2) promoter that targets the oral cavity, esophagus, and squamous forestomach^81^. These mice develop chronic inflammation that progresses to BE and EAC; progression was accelerated by exposure to bile acids. Findings show that a combination of inflammation and bile acids, the latter are components of gastroduodenal reflux that has been linked to BE→EAC progression^81, 82^. Most importantly, the bile acid-accelerated model (<u>GSE24931</u>) recapitulated the neutrophil processes that were encountered in most human datasets of EACs and GEJ-ACs (**Figure 5G**).

### Absolute neutrophil count (ANC) and neutrophil signatures prognosticate outcome in BE/EAC

We next asked how neutrophil counts and signatures impact the risk of transformation in BE and EAC outcome. First, we retrospectively analyzed data of patients with BE [NDBE, n = 72, DBE, n = 11] diagnosed between 2013 and 2017 and patients with EACs [n=30] diagnosed between 2005 and 2017, with a complete blood count within 6 months from diagnostic endoscopy (see *Methods;* **Figure 6A**). Patients were followed up for outcome until December 2021. As published before by numerous groups^83–85^, the neutrophil:lymphocyte ratio (NLR) progressively increased during NDBE→DBE→EAC progression (**Figure 6B**). However, the increase in NLR in DBE and the extent of increase in NLR in EAC were largely driven by progressive increases in the absolute neutrophil counts (ANC; **Figure 6C**) and not due to reduced absolute lymphocyte counts (ALC; **Figure 6D**). Platelet count (**Figure 6E**) and total leukocyte count (**Figure 6F**) were significantly increased in patients diagnosed with EACs; the latter is largely driven by neutrophilia and despite significant lymphopenia. ANC remained the most significant variable that tracked the risk of NDBE→DBE→EACs progression in both univariate (**Figure 6G**) and multivariate (**Figure 6H**) analyses.

**Figure 6.**
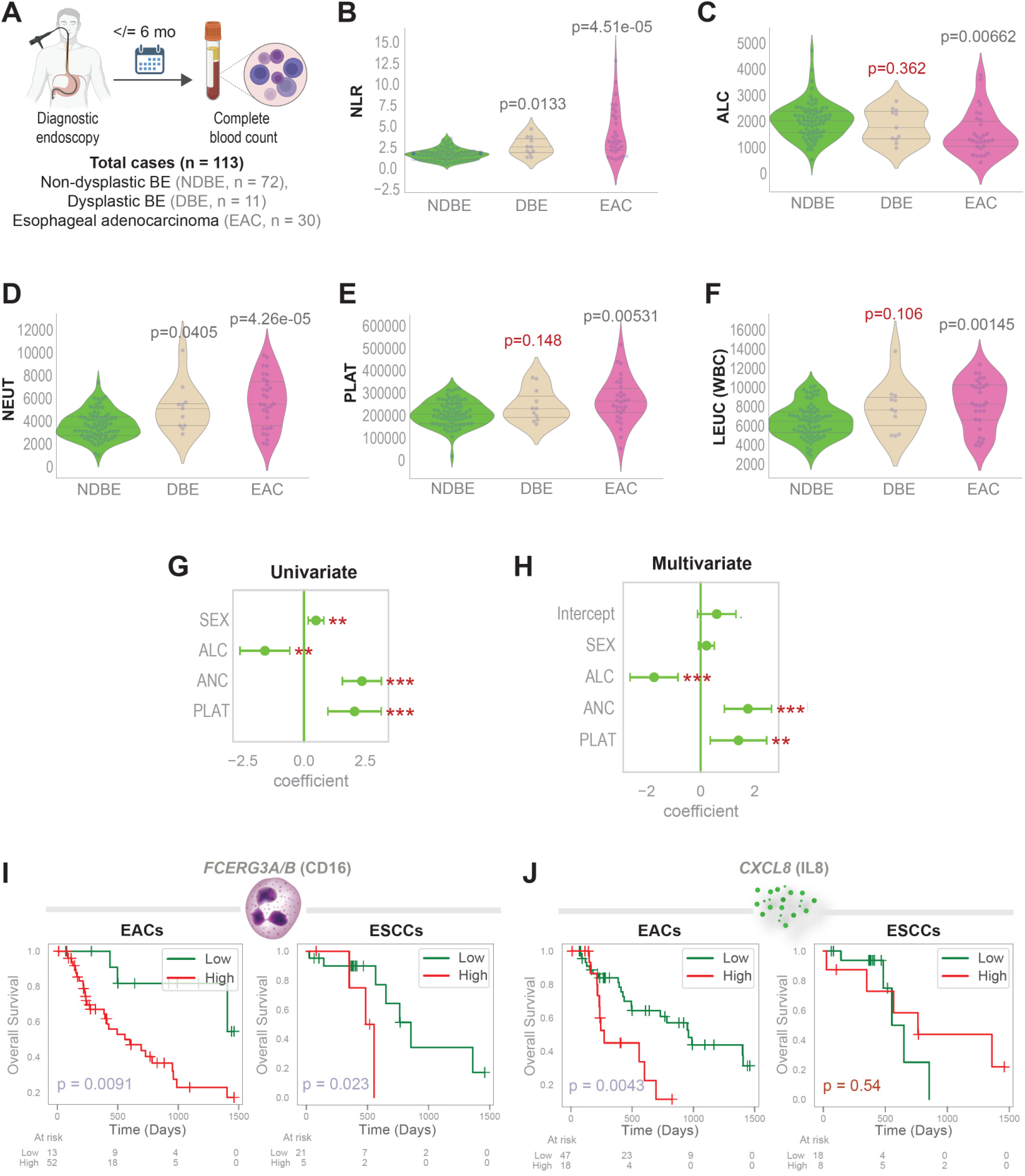
Peripheral neutrophilia and infiltrating neutrophils in tumors carry prognosticate risk of disease progression. **A.** Schematic summarizing the cross-sectional study design and cohort composition that is analyzed in panels B-H. **B-F**. Violin plots display the neutrophil:lymphocyte ratio (NLR, B), or absolute lymphocyte (ALC, C), neutrophil (NEUT, D), platelet (PLAT, E) and leukocyte (LEUK, F) counts in various patients within each diagnostic group shown in panel A. *P* values indicate comparison of each subgroup against the non-dysplastic BE (NDBE) group, as determined by Welch’s t-test. **G-H**. Univariate (G) and multivariate (H) analyses models the risk of BE to EAC progression as a linear combination of sex and the indicated hematologic parameters. Coefficient of each variable (at the center) with 95% confidence intervals (as error bars) and the *p* values were illustrated in the bar plot. The *p* value for each term tests the null hypothesis that the coefficient is equal to zero (no effect). \*\**p*≤ 0.01; \*\*\**p*≤ 0.001. **I-J**. Schematic (left) indicates the chemoattractant *CXCL8* (J) and CD16 (I; a heterodimer that is encoded by the genes *FCERG3A/B*), the marker of TAN abundance that is assessed here. Kaplan-Meier (KM) plots display the overall survival of patients with tumors stratified based on the high vs low composite scores of 2 genes (*FCERG3A, FCERG3B*) and the high vs low expression values of *CXCL8.* High and low values are obtained using StepMiner threshold +/- noise margin (See supplementary methods for detail). *p* values were determined by logrank analysis. See also **Figure S5** for KM plots on these and other signatures, analyzed in EACs, ESCCs and gastric adenocarcinomas (GCs).

The prognostic role of the EAC signature (**Figure S5A**), the neutrophil degranulation (signature derived from the EAC cluster; **Figure S5B**), the neutrophil chemoattractant, CXCL8/IL8 (**Figure S5C**), and neutrophil abundance, estimated in tumor tissues by transcripts of the marker, CD16^86^ (*FCGR3A/B*, Fc Gamma Receptor IIIa; **Figure S5D**) were also confirmed in the EAC datasets, curated from the TCGA. While the EAC and neutrophil signatures retained their prognostic impact also in ESCCs, CXCL8 did not (**Figure S5A-D**; right column). None of these signatures prognosticated outcome in gastric adenocarcinomas (GC; **Figure S5A-D**; right column). A combined CD16 and CXCL8/IL8 composite score was prognostic for EACs (**Figure 6I**), but not ESCC or GCs (**Figure 6J-K**). These results show that high ANC, high CXCL8/IL8 and high intra-tumoral neutrophil-activation signatures may all contribute as disease drivers. They also suggest that IL8-driven trafficking of neutrophils into the tissue may be a unique driver in EAC, but not in ESCC or GCs.

### Caucasians, but not African Americans mount IL8- and neutrophil-centric inflammation

The discovery of the only known race-influenced mechanism that protects cells against DNA damage, i.e., high glutathione S-transferase theta 2 (GSTT2) in AAs compared to Cau^87^ (**Figure S7A**), was made using a unique dataset containing histologically normal esophageal squamous lining-derived transcriptomics from Cau and AA subjects with normal (N) or BE diagnosis (BE) (**Figure 7A**; see **<u>Supplemental Information 1</u>**; <u>GSE77563</u>^87^). We leveraged this dataset to ask how our findings differ along the race/gender divide. We made 3 observations: **a)** the BE (**Figure 7B**-*left*) or EAC (**Figure 7B**-*right*) signatures were not different in the squamous lining of the esophagus as baseline (compare AA-N vs Cau-N); **b)** a diagnosis of BE in Cau, but not AA was associated with an induction of the EAC signatures in the histologically normal proximal squamous lining (**Figure 7B**-*right*; compare AA-BE vs Cau-BE); **c)** When these signatures, and all other signatures of tumor microenvironment and neutrophil processes were analyzed systematically, we found that the changes in gene signatures was seen in both genders (**Figure 7C-D**, compare bottom 3 rows; **Figure S6**); however, Caucasian men [Cau-BE (M)] accounted for the most significant changes in the signatures across the board. Thus, the nature of the progressive inflammation and cell states identified by our network approach is induced preferentially in Cau subjects with BE, in whom the risk of BE/EAC is ~4-5-fold higher than AAs. Finally, we found that the TIS^67, 68^ is induced in Cau subjects with BE, but not AA (**Figure 7C**), suggesting that these inhibitors may stall cell transformation in Cau subjects.

**Figure 7.**
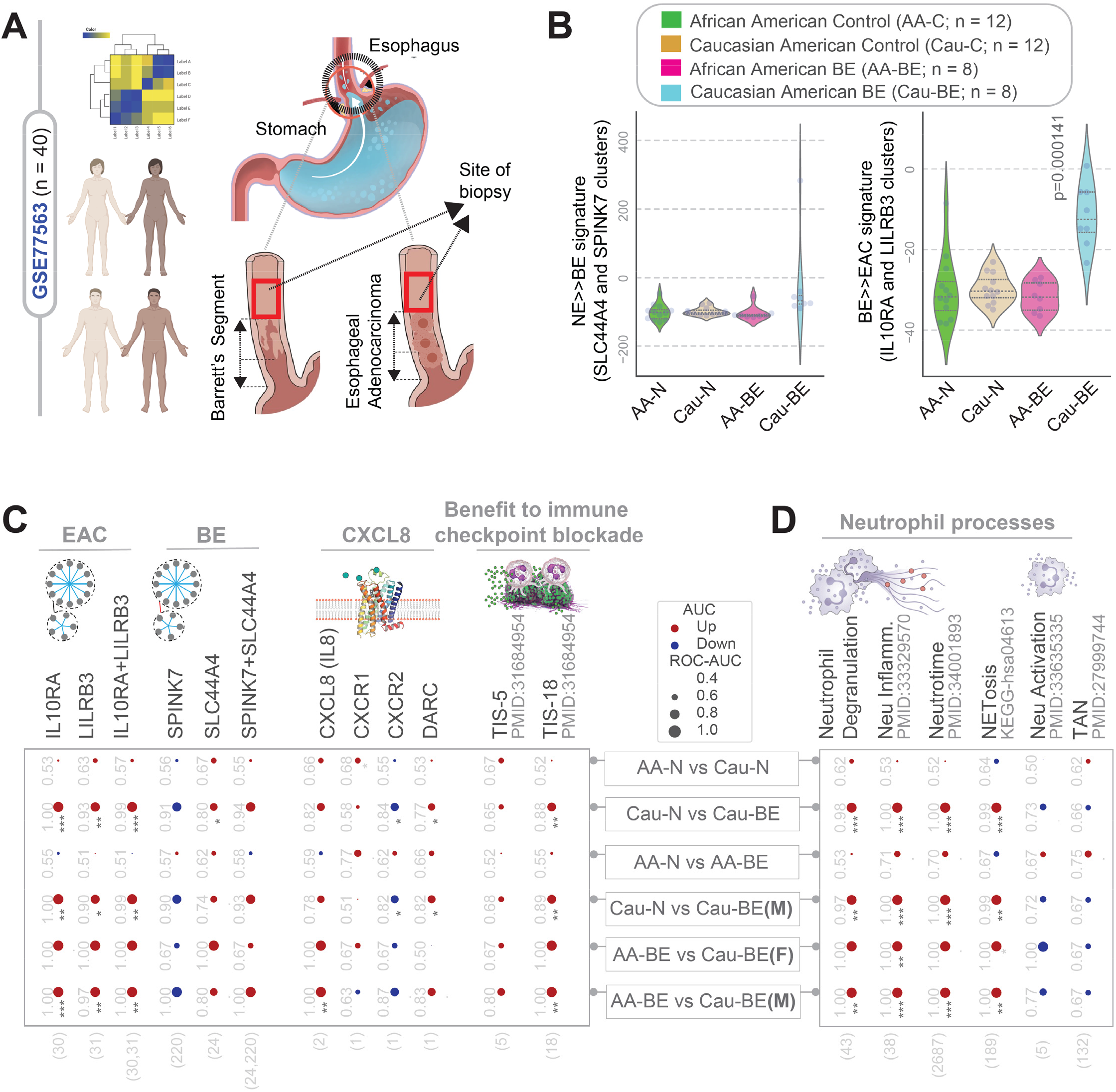
Caucasians (Cau), but not African Americans (AAs) mount IL8- and neutrophil-centric inflammation. **A.** Schematic displays the study design in <u>GSE77563</u>. Microarray studies were conducted on histologically normal squamous mucosa arising within the tubular esophagus above the gastrointestinal junction from self-identified AA or Cau subjects, who had no history of BE or EAC (normal controls), or those diagnosed with BE and/or EAC (AA-BE or Cau-BE). **B.** Violin plots showing the composite scores of upregulated gene clusters (*Left*, BE signatures; *Right*, EAC signatures) in normal esophagus from control AA or Cau subjects (AA-N and Cau-N) and esophagus from AA/Cau subjects diagnosed with BE/EACs (AA-BE and Cau-BE). *P* values indicate comparison of each sample type against the normal samples, as determined by Welch’s t-test. **C-D.** Human EAC immune microenvironment is visualized as bubble plots of ROC-AUC values (radius of circles are based on the ROC-AUC) demonstrating the direction of gene regulation (Up, red; Down, blue) for the classification of samples (gene signatures in columns; dataset and sample comparison in rows). P values based on Welch’s T-test (of composite score of gene expression values) are provided using standard code (‘‘ p> 0.1, ‘.’ p <= 0.1, * P <= 0.05, ** P <= 0.01, *** P <= 0.001) next to the ROC-AUC. Panel C displays the classification of AA vs Cau samples from control (AA/Cau-N) or BE/EAC subjects (AA/Cau-BE) in males (M) or female subjects (F) based on signatures derived from the BE>>EAC map (upregulated gene clusters *IL10RA* and *LILRB3*), NE>>BE map (downregulated *SPINK7*-cluster, and upregulated *SLC44A4*-cluster), CXCL8/IL8 and its three receptors (CXCR1/2 and DARC), and tumor inflammation signatures that predict anti-PD1 Rx (TIS-5, TIS-18) in <u>GSE77563</u>. Panel D displays the classification of same samples in C based on neutrophil signatures (NETosis, Degranulation, Inflammation, Neutrotime) in <u>GSE77563</u>. Violin plots for selected neutrophil signatures in AA-BE vs Cau-BE samples are displayed in **Figure S6**.

It is noteworthy that the protective high levels of expression of GSTT2 in AAs over Cau^87^ (see legend, **Figure S7A**) is observed both in healthy controls as well as in subjects with BE/EAC (**Figure S7B**). GSTT2 expression inversely correlated with the EAC signature (**Figure S7C**) and its subset of neutrophil degranulation signature (**Figure S7D**). These findings indicate that the neutrophil/IL8 centric inflammation that is mounted exclusively during BE→EAC progression in the Cau subjects (**Figure 7B**-*right*) occurs in the vulnerable setting of low GSTT2 in the same subjects. Because GSTT2 protects against DNA damage, findings suggest that racial differences in GSTT2 may protect AAs from or predispose Cau to more DNA damage in the setting of the IL8-neutrophil centric inflammation.

### SNPs that increase or decrease ANC are oppositely enriched during BE→EAC progression

We next asked if race and/or ethnicity may intersect directly with ANC and the risk of BE→EAC progression. Populations of African descent are known to have low ANCs compared to others^88^ while those of Hispanic/Latino (HL) descent have high ANCs compared to non-Hispanic white (NHW^89^). Single nucleotide polymorphisms (SNPs) that either increase (such as in the US Hispanic/Latino population^90^) or decrease (such as in the US AA population^91, 92^) ANCs have been identified. In the case of AAs, a study^92^ that pooled >6,000 AA samples from 3 studies [the Jackson Heart Study (JHS), the Atherosclerotic Risk in Communities (ARIC) Study, and the Health, Aging and Body Composition (Health ABC) Study], the homozygous SNP (rs2814778, which disrupts a binding site for the GATA1 erythroid transcription factor, resulting in a ACKR1-null phenotype (**Figure 8A**) was found to be predictive of low ANC in AAs above beyond the previously described admixture association. This study established a *bona fide* phenotype (low ANC) for this genetic variant and demonstrated that the causal variant must be at least 91% different in frequency between AA vs Cau. In the absence of a longitudinal study that monitors BE progression in AA vs Cau subjects, we leveraged a case-control study^93^ that included 80 patients with BE (40 who progressed to EACs and 40 who did not, i.e., non-progressors); this cohort was selected from a larger case-cohort study within the Seattle Barrett’s Esophagus Program (SBEP) at the Fred Hutchinson Cancer Research Center^94^. Germline data from this cohort was analyzed for the occurrence of 10 SNPs that increase or decrease the ANC, as determined in various studies in the US (see complete list in **<u>Supplemental Information 6</u>**). All 3 genes (*DARC/ACKR1, ABCC1* and *HMMR*), impacted by 4 of the 6 tested risk alleles, including *rs2814778* [the allele maximal risk, across studies, and the one that confers the risk of benign ethnic neutropenia (BEN^*95*^)] were significantly enriched among non-progressors, whereas 3 of the 4 protective alleles were significantly enriched among the progressors (**Figure 8B**). Among other SNPs that are associated with drug induced neutropenia in the non-US population, only one was significant (i.e., *CYP39A1*; **<u>Supplemental Information 6</u>**). The opposing patterns of enrichment and de-enrichment of neutropenia-protective and risk alleles, respectively, among BE→EAC progressors was significant (Welch’s t-test *p* = 0.03978; **Figure 8B**). As expected, somatic mutations in genes within the EAC clusters (**Figure S7A**) or on genes associated with neutrophil function/number (**Figure S7B**) were more frequent among progressors compared to non-progressors, and the frequency increased further in tumors with high mutation burden.

**Figure 8.**
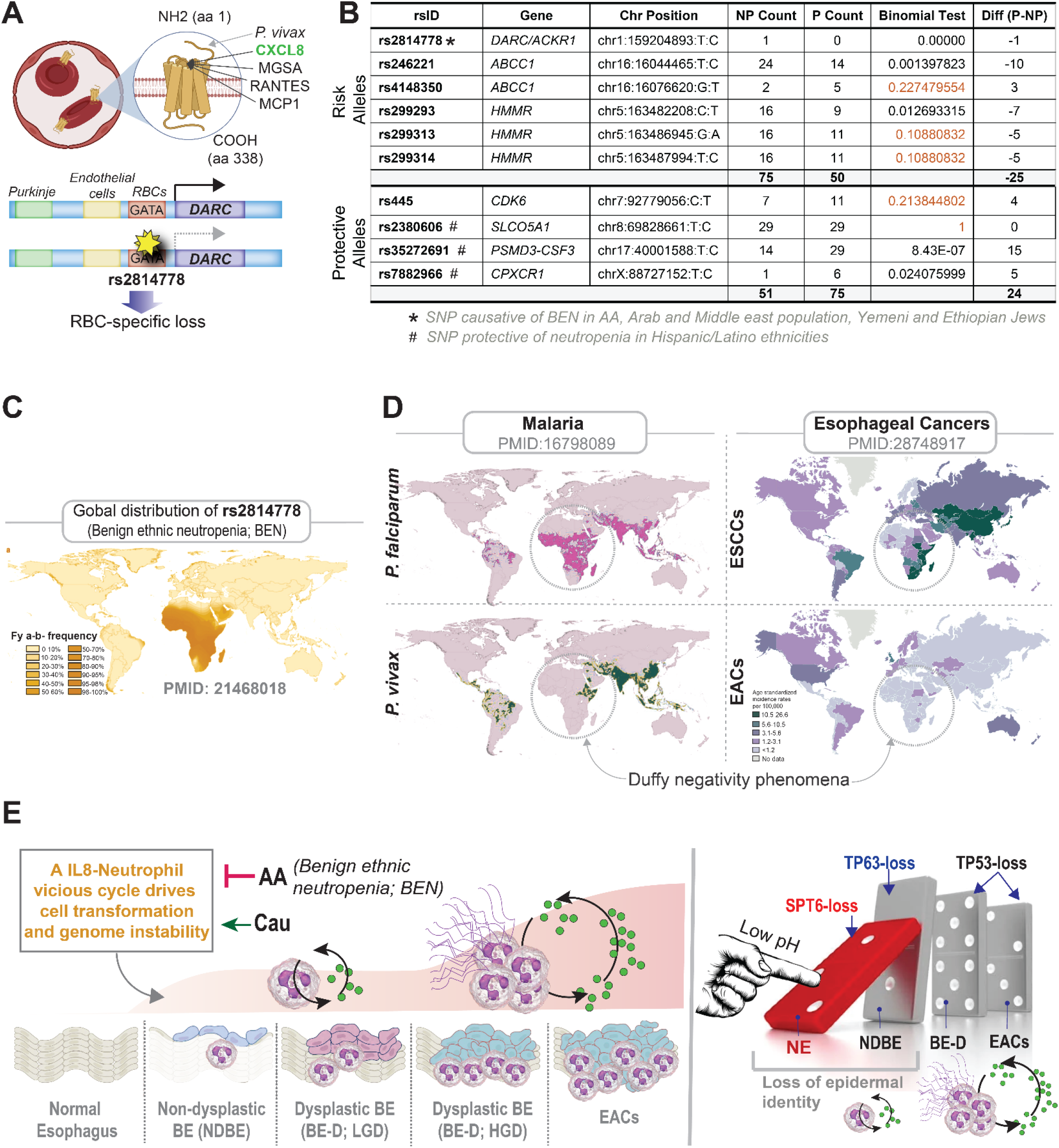
Ethnic neutropenia may reduce the risk of transformation in BE. **A**. Schematic summarizing the ligands that bind RBC-localized *DARC/ACKR1* scavenger (A) and the impact of the African polymorphism on RBC-specific loss of *DARC*. **B.** Table showing the frequency of neutropenia risk alleles (top) or protective alleles (bottom) among patients with BE (Seattle BE program; n = 80) who progress (Prog, P; n = 40) or not (Non-prog, NP; n = 40) to EAC. Statistical significance was determined for each SNP using Fisher’s test with binomial probability distribution. See also **Supplementary Information 6**. **C**. Global prevalence of the African *Duffy-null* polymorphism that causes benign ethnic neutropenia. **D**. The prevalence of malaria (left) and the age-adjusted incidence of ECs (right) are displayed side by side. Black interrupted circles highlight how inability of *P. vivax* to attach to RBCs in *Duffy*-null zones (see C) protect Africans from that specific form of malaria (bottom-left) but not from *P. falciparum* (top-left). Circles on highlight similar differences in ESCC vs EAC in the African population. **E.** Summary and working model: *<u>Left</u>:* A vicious IL8↔neutrophilic storm, perhaps biphasic, may be critical for driving the metaplasia-dysplasia cascade during NE→BE→EAC progression. Due to the AA *Duffy-null* polymorphism which manifests as benign ethnic neutropenia, some races/ethnicities (like AA and Hispanic/Latino) are protected from both the biphasic IL8/neutrophil-centric inflammatory storm *<u>and</u>* the metaplasia-dysplasia cascade. *<u>Right</u>:* As for epithelium intrinsic events, we identified a molecular mechanism that permits NE→BE transition. This mechanism is triggered by acid exposure (at least in part), suppresses the expression of SPT6, which in turn stalls TP63 function and expression, and a resultant loss in keratinocyte cell fate and gain in metaplastic features. These epithelium-intrinsic mechanisms are likely to be fuelled by the vicious IL8↔neutrophilic storm pinpointed by our computational models.

### Ethnic neutropenia may reduce EAC risk

We hypothesized that if the IL8↔neutrophil axis drives cell transformation and hence, the risk of EACs, then ethnic prevalence of polymorphisms that impact either IL8 signaling and/or neutrophil abundance might influence such risk. We found that ethnicities with the lowest incidence of BE/EAC (i.e., AAs), also have BEN^*95*^, the most common form of neutropenia worldwide. In BEN, a homozygous SNP (*rs2814778*) impacts the functions of *DARC* (*only in RBCs*), which encodes is a 7-TM receptor^96^ that selectively scavenges “inflammatory” chemokines e.g., IL8 and *CCL5*, both of which enhance neutrophil recruitment^97–99^ (**Figure 8A**). The AA *Duffy* polymorphism has explained many mysterious racial disparities in modern medicine, e.g., high susceptibility to HIV^91^, but protection against *P. vivax* malaria^92^, high sickle cell-related complications, aging phenotypes^100^, and acute lung injury^101^ among AAs. Its impact on neutrophil count is generalizable also to Yemenite Jews^102^ and Hispanics/Latinos^90^. We looked for the ‘*Duffy* negativity phenomenon’ which was first described in malaria^103^; this phenomenon refers to the geographic distribution of the *Duffy* negative genotype, *Fy^a-/b-^*, predominantly in sub-Saharan Africa (≥95% Duffy negativity frequency; CI 75%-95%; **Figure 8C**) which coincides with the phenotype of near complete protection from *P. vivax* in the same regions (**Figure 8D**-*bottom panel*). It is noteworthy that *Duffy* negativity does not offer protection from *P. falciparum* (**Figure 8D**-*top panel*). A strikingly similar contrasting pattern was seen when we compared the global age-adjusted incidence rates of ESCCs and EACs (**Figure 8E**); the African and Saudi Arabian regions the *Duffy* negative genotype, *Fy^a-/b^* (**Figure 8C**) showed low incidence of EACs, but moderate to high incidence of ESCCs. These findings suggest that BEN could offer selective protection from EACs (just as it does for *P. vivax*) in individuals of African descent. Findings also suggest that BEN, which is widely prevalent in other ethnicities (e.g., Africans^104–106^, AAs, Arabs^107^, Yemenite and black Ethiopian Jews^105^, and to a lesser extent also in Latinos^90^) but < 1% in the non-Hispanic white population in the US^89^ is a possible risk modifier (protective) for BE→EAC progression.

## CONCLUSIONS

The major discovery we report here, using an off-the-beaten path network transcriptomic approach, are insights into the cellular continuum states during the metaplasia→dysplasia→neoplasia cascade in adenocarcinomas of the esophagus and gastroesophageal junction. Findings enable us to draw four major conclusions, some with immediate and impactful translational relevance.

### All EACs must originate from BE

First, our findings support a long-suspected tenet, originally proposed in 2000^108^” i.e., that all EACs arise in BE. That tenet was substantiated *via* recent multiscale computational modeling studies^109^, through single-cell genomics and lineage tracking studies^110^, and now through a Boolean logic-based model (current work). These three approaches independently verify that BE is the invariant precursor to EACs. The same Boolean logic-based model confirmed that, unlike EACs, ESCCs do not transition through metaplastic BE states (as expected). TP63 and SPT6 were suppressed in the squamous lining of the esophagus proximal to the BE segment (in **Figure 3D-E**), and BE/EAC signatures and neutrophil inflammatory milieu was similarly observed in the normal squamous lining of the esophagus proximal to BE/EAC lesions (in **Figure 7A-B**) suggests that the histologically so-called ‘normal’ esophageal lining is abnormal by all molecular (protein and gene expression) metrics among patients with BE/EAC. This evidence lends support to a transcommitted esophageal keratinocyte being a cell of origin of BE and EAC/GEJ-ACs, as has been suggested by several independent groups^44–47^.

### Neutrophilic assault assists cell transformation

Second, we pinpointed a 2-step waveform of an IL8-neutrophil centric tumor immune microenvironment that is increased first during metaplastic transition and more prominently later during neoplastic transformation (see **Figure 8F** legend). This biphasic-wave pattern may explain why long-standing BE carries low risk of EAC^111^, presumably because a 2^nd^ wave/’hit’ of immune storm is required for cell transformation. Induction of IL8 and its receptors, CXCR1/2^49, 112–116^, increased presence of neutrophils in BE tissues^117^ and circulation^83, 84^ have been reported and increased NETosis within EACs in the setting of peripheral leukocytosis has been shown to carry worse prognosis^118^. In showing that the IL8-neutrophil centric tumor microenvironment is prominently induced in Cau, but not in AAs, which mirrors the ~4-5 fold higher risk of EACs in Cau compared to AAs^119^, our findings suggest that this inflammatory microenvironment is likely to be a driver event. Besides the tumor environment, we showed through multivariate analyses that a high ANC in circulation is an independent determinant of NDBE→DBE→EAC progression. We provide epidemiologic evidence and propose that ethnic neutropenia is a possible risk modifier (protective) for BE→EAC progression in AAs, Arabs, and other ethnicities. The fact that peripheral neutrophilia and intratumoral signatures of neutrophil processes are aligned, i.e., both carry poor prognosis in EACs is not unusual because prior studies have confirmed that both peripheral neutrophilia and the infiltration of TANs are associated with worse outcome in diverse cancers^120^. Although not investigated here, the insights into the IL8↔neutrophil centric immune microenvironment aid in formulating testable hypotheses on disease pathogenesis and prioritize a class of neutrophil-targeted therapeutics. For example, TAN infiltration may accelerate adenocarcinomas by impacting a slew of pro-cancer processes (reviewed in^121–123^), and hence, anti-neutrophil drugs could prove beneficial as both single agent as well as adjuvant therapy. Because neutrophil infiltration in tumor is associated with key features of resistance to immune checkpoint inhibitors^124^, and high neutrophil counts with high tumor mutation burden in diverse cancers (including esophageal) is known to reduce the efficacy of the checkpoint inhibitors^125^, our results predict that neutrophil-targeted therapeutics will synergize with checkpoint inhibitors.

### *Ethnic disparity in EACs/GEJ-ACs* may stem from *DARC polymorphism*

Third, our studies shed valuable insights into how *DARC* polymorphism *rs2814778* may shape the risk of EACs. For example, neutrophil processes are high in both EACs and ESCCs, and yet, the existence of a *Duffy* negativity phenomenon in EACs (that the *Fy^a-b-^* genotype is associated with low incidence of EACs, but not ESCCs) suggests that the *Duffy* polymorphism impacts EACs through mechanisms other than being the most important genetic determinant of BEN. *DARC* polymorphism *rs2814778* is known to impact serum levels of MPO^126, 127^ and IL8^128^ and their coexistence with IL8 polymorphisms is known to impact the prevalence of periodontitis in certain ethnicities^129^. Because EACs, but not ESCCs, significantly induce *CXCL8,* it is possible that the infiltration of TANs in EACs is gradient driven, and that gradient is maintained by the RBC-localized scavenger of *CXCL8, DARC* (serving as a cytokine ‘sink’^130–132^). We cannot rule out that *DARC*-mediated chemokine sequestration may modulate tumor microenvironment, angiogenesis and metastasis, which has been described in other cancers (reviewed in^91^). As for the observed gender disparity (higher risk of EACs/GEJ-ACs in men), a more reactive neutrophil state^133^ or prevalence of autoimmune or idiopathic neutropenia^89, 134^ may offer some explanation, but ethnic neutropenia due to *DARC* polymorphism *rs2814778* is just as important of a determinant of low ANC in females as in men^135^.

### EACs and GEJ-ACs behave similarly

Fourth, by showing that EACs and GEJ-ACs share similar gene signatures and tumor microenvironments, our study objectively establishes the degree of similarity between the two tumors, both with rapidly rising incidence, at a fundamental molecular level. This finding is in keeping with the fact that, much like EACs, GEJACs are also associated with short and long segments of BE suggesting that they arise from underlying metaplastic epithelium^136^. It is also consistent with the fact that the entity of familial BE has been seen in ~7.3% of patients presenting with BE, EACs, or GEJACs^137^. It is possible that much like EACs, GEJ-ACs evolve through the intestinal metaplastic continuum and that neutrophil-targeted or other immune checkpoint therapies that emerge in EACs are expected to have crossover benefits in GEJ-ACs.

## STUDY LIMITATIONS

The biggest limitation of this study and our approach is the availability of data (or lack thereof). For example, due to lack of datasets with normal esophagus, gastric cardia, BE and EAC/GEJ-ACs all in one cohort, the logical model of EAC and GEJ-AC initiation we cannot rule out that cardia→BE→EAC is another plausible path. ANC, serum, and tumor IL8 expression, TAN and tumor neutrophil signatures were not evaluated in the same cohort; well-curated prospective studies will be necessary to connect these dots - independently and as composite measures -- with outcome. Such studies will help understand the role of IL8-gradient (serum to tumor), neutrophilia and TAN infiltration. Although this study proposes BEN as a modifier of EAC risk, the impact of BEN on BE→EAC progression requires a study that is controlled for all variables (including race, smoking, BMI, etc) and genotyped (for *DARC* polymorphism rs2814778) and phenotyped (monitored for ANC and BE/EAC progression). Because smokers in all ethnic groups had a higher neutrophil count than non-smokers^89^ (greatest effect on increments of ANC among Cau), it is possible that neutrophil processes and neutrophilia intersects with the established smoking-related risk of EACs and GEJ-ACs^138^; this was not investigated here. This study also did not address the mechanism(s) by which peripheral neutrophils/TANs may accelerate adenocarcinomas in the esophagus and the GE-junction and how *DARC* polymorphism *rs2814778* may offer protection. Tumor-neutrophil co-culture studies incorporating IL8 gradients and titrated neutrophil counts may yield valuable insights into how neutrophil infiltration impact tumor mutation burden.

## METHODS

Detailed methods for computational modeling, AI-guided prediction and validation and description of validation models are presented in Supplementary Online Materials and mentioned in brief here.

### Computational Approach

An overview of the key approaches is shown in **Figure 1**. Modeling continuum states within the metaplasia→dysplasia→neoplasia cascade was performed using <u>Bo</u>olean <u>N</u>etwork <u>E</u>xplorer (*BoNE*)^29^. We created an asymmetric gene expression network, first for metaplastic progression from normal esophagus to BE, and separately, for the dysplastic→neoplastic cascade during BE to EAC progression, using a computational method based on Boolean logic ^9, 10^, ^139^. To build the BE/EAC network, we analyzed two publicly available colon-derived transcriptomic datasets [<u>GSE100843</u>^30^ and <u>GSE39491</u>^31^ for BE and <u>E-MTAB-4045</u> for EACs (see **<u>Supplemental Information 1</u>**). These two datasets (‘test cohorts’) were independently analyzed and the resultant signatures were kept separate from each other at all times. A <u>Bo</u>olean <u>N</u>etwork <u>E</u>xplorer (*BoNE*; see Supplementary Methods) computational tool was introduced, which uses asymmetric properties of Boolean implication relationships (BIRs as in MIDReG algorithm^9^) to model natural progressive time-series changes in major cellular compartments that initiate, propagate and perpetuate cellular state change and are likely to be important for BE/EAC progression. *BoNE* provides an integrated platform for the construction, visualization and querying of a network of progressive changes much like a disease map (in this case, BE and EAC-maps) in three steps: First, the expression levels of all genes in these datasets were converted to binary values (high or low) using the *StepMiner* algorithm^140^. Second, gene expression relationships between pairs of genes were classified into one-of-six possible BIRs and expressed as Boolean implication statements; two symmetric Boolean implications “equivalent” and “opposite” are discovered when two diagonally opposite sparse quadrants are identified and four asymmetric relationships, each corresponding to one sparse quadrant. While conventional symmetric analysis of transcriptomic datasets can recognize the latter 2 relationships, such approach ignores the former. BooleanNet statistics is used to assess the significance of the Boolean implication relationships^9^. Prior work^28^ has revealed how the Boolean approach offers a distinct advantage from currently used conventional computational methods that rely exclusively on symmetric linear relationships from gene expression data, e.g., Differential, correlation-network, coexpression-network, mutual information-network, and the Bayesian approach. The other advantage of using BIRs is that they are robust to the noise of sample heterogeneity (i.e., healthy, diseased, genotypic, phenotypic, ethnic, interventions, disease severity) and every sample follows the same mathematical equation, and hence is likely to be reproducible in independent validation datasets. The heterogeneity of samples in each of the datasets used in this study is highlighted in **<u>Supplemental Information 1</u>**. Third, genes with similar expression architectures, determined by sharing at least half of the equivalences among gene pairs, were grouped into clusters and organized into a network by determining the overwhelming Boolean relationships observed between any two clusters ^9, 10^. In the resultant Boolean implication network, clusters of genes are the nodes, and the BIR between the clusters are the directed edges; *BoNE* enables their discovery in an unsupervised way while remaining agnostic to the sample type. All gene expression datasets were visualized using Hierarchical Exploration of Gene Expression Microarrays Online (HEGEMON) framework^141^.

### Human subjects

For assessing the impact of absolute neutrophil count (ANC) on BE/EAC diagnosis and outcome of EACs, we retrospectively analyzed patients with biopsies reporting BE between 2013 and 2017 and with a complete blood count within 6 months from the endoscopy, as well as patients with esophageal adenocarcinoma (EAC) at a tertiary care center in Brazil (Hospital de Clínicas dePorto Alegre). Cases (n = 113) were classified as non-dysplastic BE (NDBE, n = 72), dysplastic BE (DBE, n = 11) and EAC (n = 30)^83^. Data collected from endoscopic histopathological reports were histologic diagnosis and adherence to Seattle Protocol^142, 143^. The study was approved by the Brazilian National Committee on Research Ethics (CONEP), registered by number CAAE-81068617.2.0000.5327.

For collecting esophageal biopsies for IHC we enrolled patients undergoing endoscopies as a part of their routine care and follow-up at UC San Diego’s Center for Esophageal Diseases. Patients were recruited and consented using a study proposal (Project ID#200047) approved by the Human Research Protection Program (HRPP) Institutional Review Board of the University of California, San Diego (Project ID#200047).

Human keratinocyte-derived BE organoids were created^39^ and validated^144^ previously using an approved IRB (Project ID # 190105) that covers human subject research at the UC San Diego HUMANOID Center of Research Excellence (CoRE). For all the deidentified human subjects, information including age, gender, and previous history of the disease, was collected from the chart following the rules of HIPAA. The study design and the use of human study participants was conducted in accordance with the criteria set by the Declaration of Helsinki.

## ACKNOWLEDGEMENTS

This work was supported by NIH AI141630 and CA100768 (to P.G), GM138385 (to D.S), T32 GM8806 (to D.V) and AI155696, UG3TR003355 and UG3TR002968 (to D.S, P.G and S.D), P30 CA023100 (KC and CG), and K23 DK125266 (RY). P.G, D.S, R.Y and S.D were also supported by research gift funds from the Torey Pines Foundation. KC and CG were also supported by a UCSD Academic Senate Grant (RG103468). We thank Donald Pizzo (UCSD Pathology Histologic Biomarkers Core), Madeline Gerytak and Vanessa Castillo (UCSD) for technical and logistical support.

## AUTHOR CONTRIBUTIONS TO MANUSCRIPT

DS and PG conceptualized the project; DS and DV, under the supervision of PG and DS carried out computational analyses; DS contributed all software used in this work; RY provided access to patients and secured all tissue samples used in this study; VGH carried out the immunohistochemical studies under the supervision of PG; CT and SD were responsible for the BE organoid model studies; CG, PH and KC carried out the mutation and SNP analyses in BE datasets, accessed through the pre-cancer genome atlas (PCGA) pipeline co-created by LA and SML. DS and PG prepared figures for data visualization and wrote the original draft of the manuscript. All authors provided input and edited and revised the manuscript. All co-authors approved the final version of the manuscript. PG coordinated and supervised all parts of the project and administered the project.

## DATA AVAILABILITY

All data is available in the main text or the supplementary materials. Publicly available datasets used are enlisted in **<u>Supplemental Information 1</u>**. The software codes are publicly available at the following links: https://github.com/sahoo00/BoNE and https://github.com/sahoo00/Hegemon.

## SUPPLEMENTAL INFORMATION (Excel datasets)

**<u>Supplemental Information 1</u>:** Excel sheet with list of all transcriptomic datasets used in this work.

**<u>Supplemental Information 2</u>:** Excel sheet with list of BE-map derived genes and reactome pathways.

**<u>Supplemental Information 3</u>:** Excel sheet with list of EAC-map derived genes and reactome pathways.

**<u>Supplemental Information 4</u>:** Excel sheet with list of all neutrophil process-related signatures.

**<u>Supplemental Information 5</u>:** Excel sheet with hematologic parameters assessed in a cross-sectional retrospective study on 113 BE and EAC patients.

**<u>Supplemental Information 6</u>:** Excel sheet with SNP frequency among progressors and non-progressors in a case-control genomics study [the Seattle Barrett’s Esophagus Program (SBEP)].

## Abbreviations

BE: Barrett’s metaplasia of the esophagus
EAC: Esophageal adenocarcinoma
GEJ: Gastroesophageal junctionx
Cau: Caucasian
AA: African American
SNP: single nucleotide polymorphism
*DARC*: Duffy antigen/ receptor for chemokines

