## Supplementary Online Materials for "AI-assisted Discovery of an Ethnicity-influenced Driver of Cell Transformation in Esophageal and Gastroesophageal Junction Adenocarcinomas"

**Affiliations**: [1]Department of Cellular and Molecular Medicine, University of California San Diego; [2] Department of Medicine, University of California San Diego; [3] HUMANOID Center of Research Excellence (CoRE), University of California San Diego; [4] Moore's Comprehensive Cancer Center, University of California San Diego; [5] Department of Gastrointestinal Surgery, Hospital de Clínicas de Porto Alegre, 2350 Ramiro Barcellos Street, Porto Alegre, RS, 90035-003; [6] Department of Pediatrics, University of California San Diego; [7] Division of Biomedical Informatics, University of California San Diego; [8] Postgraduate Program in Medicine; Surgical Sciences, Federal University of Rio Grande do Sul, 2400 Ramiro Barcellos Street, Porto Alegre, RS, 90035-003; [9] Medical School of Federal University of Rio Grande do Sul 2400 Ramiro Barcellos Street, Porto Alegre, RS, 90035-003; [10] Department of Pathology, University of California San Diego; [11] Department of Computer Science and Engineering, Jacob's School of Engineering, University of California San Diego.

†Equal contributions.

### *CORRESPONDING AUTHOR CONTACT INFORMATION

**Debashis Sahoo, Ph.D**.; Associate Professor, Department of Pediatrics, University of California San Diego; 9500 Gilman Drive, MC 0703, Leichtag Building 132; La Jolla, CA 92093-0703. Phone: 858-246-1803: Fax: 858-246-0019:

**Vinicius Campos, M.D.;** Department of Gastrointestinal Surgery, Hospital de Clínicas de Porto Alegre, 2350 Ramiro Barcellos Street, Porto Alegre, RS, 90035-003.

**Rena Yadlapati**, **M.D**.; Associate Professor, Division of Gastroenterology; Department of Medicine, University of California San Diego; 9500 Gilman Drive, MC 0703, La Jolla, CA 92093-0887.

**Kathleen Curtius**, **Ph.D**.; Assistant Professor, Division of Biomedical Informatics, Department of Medicine, University of California San Diego; 9500 Gilman Drive, MC 0703, Biomedical Research Facility-2; La Jolla, CA 92093-0728. Phone: 619-777-3296:

**Pradipta Ghosh, M.D**.; Professor, Departments of Medicine and Cell and Molecular Medicine, University of California San Diego; 9500 Gilman Drive (MC 0651), George E. Palade Bldg, Rm 232; La Jolla, CA 92093. Phone: 858-822-7633:

**This PDF file includes:**

- **Materials and Methods**
- Supplementary Text- - **Not Applicable**
- **Supplementary Figures. S1 to S8**
- **Supplementary Tables 0** (separately uploaded)
- Captions for Movies – **Not Applicable**
- Captions for Audio - **Not Applicable**
- Captions for External Data - **Not Applicable**

### Materials and Methods

#### Computational Approaches

*Barrett's and Esophageal Adenocarcinoma datasets used for network analysis*

One microarray dataset (GSE100843; n = 76, 36 Normal esophageal squamous mucosa and 40 Barrett's esophagus segment) is used to perform Boolean Implication network analysis of Barrett's esophagus (BE) samples, and another microarray dataset (GSE39491, n = 80, 40 normal, 40 BE) is used to train a network model to distinguish normal vs BE samples. Boolean Implication Network analysis of the Esophageal Adenocarcinoma (EAC) is performed using RNA-seq dataset (E-MTAB-4054, n = 63, 19 normal, 19 BE without dysplasia, 8 BE with low-grade dysplasia, 17 EAC). All training and validation dataset (**Supplementary Information 1**) were downloaded from National Center for Biotechnology Information (NCBI) Gene Expression Omnibus website (GEO)[1-3] or European Molecular Biology Laboratory (EMBL) European Bioinformatics Institute (EMBL-EBI) ArrayExpress website[4]. All gene expression datasets (**Supplementary Information 1**) were processed separately using the Hegemon data analysis framework [5-7]. We did not combine datasets that belong to two different platforms. See **Supplemental Information 1** for the degree of heterogeneity among samples in the datasets used in this work.

*Test cohort selection*

Two different test cohorts were used to build the network and perform machine learning for BE: GSE100843 and GSE39491. Both GSE100843 and GSE39491 are microarray datasets that included reasonable number (>= 30) of normal esophageal squamous mucosa and reasonable number (>= 40) of BE samples. Since the number of samples in these cohort are less than 100, which is on the lower side for comprehensive Boolean analysis, network is built on GSE100843 (n = 76) and machine learning is performed on an independent dataset GSE39491 (n = 80) to cover the entire spectrum of gene expression dataset from different microarray platforms. Surprisingly, variation in gene expression were good enough to use our standard BooleanNet statistic ($S > 3$ and $p < 0.1$) to identify Boolean Implication relationships with n = 76. Only one cohort was used to build the network and perform machine learning for EAC: E-MTAB-4054. E-MTAB-4054 (n = 63) is the only large RNASeq dataset available that provided high-quality measurements of mRNA extracted from normal, BE and EAC tissue samples. Since all these cohorts have small number of samples, reliability of Boolean analysis is low, and the results need to be supported by large and strong groups of validation datasets.

*Boolean Analysis*

*Boolean logic* is a simple mathematic relationship of two values, i.e., high/low, 1/0, or positive/negative. The Boolean analysis of gene expression data requires first the conversion of expression levels into two possible values. The *StepMiner* algorithm is reused to perform Boolean analysis of gene expression data [8]. The *Boolean analysis* is a statistical approach which creates binary logical inferences that explain the relationships between phenomena. Boolean analysis is performed to determine the relationship between the expression levels of pairs of genes. The *StepMiner* algorithm is applied to gene expression levels to convert them into Boolean values (high and low). In this algorithm, first the expression values are sorted from low to high and a rising step function is fitted to the series to identify the threshold. Middle of the step is used as the *StepMiner* threshold. This threshold is used to convert gene expression values into Boolean values. A noise margin of 2-fold change is applied around the threshold to determine intermediate values, and these values are ignored during Boolean analysis. In a scatter plot, there are four possible quadrants based on Boolean values: (low, low), (low, high), (high, low), (high, high).

*Invariant Boolean implication relationships*

A Boolean implication relationship is observed if any one of the four possible quadrants or two diagonally opposite quadrants are sparsely populated. Based on this rule, there are six different kinds of Boolean implication relationships. Two of them are symmetric: equivalent (corresponding to the highly positively correlated genes), opposite (corresponding to the highly negatively correlated genes). Four of the Boolean relationships are asymmetric, and each corresponds to one sparse quadrant: (low => low), (high => low), (low => high), (high => high). BooleanNet statistics (Equations listed below) is used to assess the sparsity of a quadrant and the significance of the Boolean implication relationships [8, 9]. Given a pair of genes A and B, four quadrants are identified by using the StepMiner thresholds on A and B by ignoring the Intermediate values defined by the noise margin of 2-fold change (+/- 0.5 around StepMiner threshold). Number of samples in each quadrant are defined as $a_{00}$, $a_{01}$, $a_{10}$, and $a_{11}$. Total number of samples where gene expression values for A and B are low is computed using following equations.

$nA_{low} = (a_{00} + a_{01}), nB_{low} = (a_{00} + a_{10}),$

Total number of samples considered is computed using following equation.

$total = a_{00} + a_{01} + a_{10} + a_{11}$

Expected number of samples in each quadrant is computed by assuming independence between A and B. For example, expected number of samples in the bottom left quadrant $e_{00} = \hat{n}$ is computed as probability of A low (($a_{00}$ + $a_{01}$)/total) multiplied by probability of B low (($a_{00}$ + $a_{10}$)/total) multiplied by total number of samples. Following equation is used to compute the expected number of samples.

$n = a_{ij}, \hat{n} = (nA_{low}/total * nB_{low}/total) * total$

To check whether a quadrant is sparse, a statistical test for ($e_{00} > a_{00}$) or ($\hat{n} > n$) is performed by computing $S_{00}$ and $p_{00}$ using following equations. A quadrant is considered sparse if $S_{00}$ is high ($\hat{n} > n$) and $p_{00}$ is small.

$$S_{ij} = \frac{\hat{n} - n}{\sqrt{\hat{n}}}$$

$$p_{00} = \frac{1}{2}\left(\frac{a_{00}}{(a_{00} + a_{01})} + \frac{a_{00}}{(a_{00} + a_{10})}\right)$$

A threshold of $S_{00}$ > sthr and $p_{00}$ < pthr to check sparse quadrant. A Boolean implication relationship is identified when a sparse quadrant is discovered using following equation.

***Boolean Implication*** = ($S_{ij}$ > sthr, $p_{ij}$ < pthr)

A relationship is called Boolean equivalent if top-left and bottom-right quadrants are sparse.

*Equivalent* = $(S_{01} > sthr, \ P_{01} < pthr, S_{10} > sthr, P_{10} < pthr)$

Boolean opposite relationships have sparse top-right ($a_{11}$) and bottom-left ($a_{00}$) quadrants.

$$Opposite = (S_{00} > sthr, \ P_{00} < pthr, S_{11} > sthr, P_{11} < pthr)$$

Boolean equivalent and opposite are symmetric relationship because the relationship from A to B is same as from B to A. Asymmetric relationship forms when there is only one quadrant sparse (A low => B low: top-left; A low => B high: bottom-left; A high=> B high: bottom-right; A high => B low: top-right). These relationships are asymmetric because the relationship from A to B is different from B to A. For example, A low => B low and B low => A low are two different relationships.

A low => B high is discovered if bottom-left ($a_{00}$) quadrant is sparse and this relationship satisfies following conditions.

*A low => B high* = $(S_{00} > sthr, \ P_{00} < pthr)$

Similarly, A low => B low is identified if top-left ($a_{01}$) quadrant is sparse.

*A low => B low* = $(S_{01} > sthr, \ P_{01} < pthr)$

A high => B high Boolean implication is established if bottom-right ($a_{10}$) quadrant is sparse as described below.

*A high => B high* = $(S_{10} > sthr, \ P_{10} < pthr)$

Boolean implication A high => B low is found if top-right ($a_{11}$) quadrant is sparse using following equation.

*A high => B low* = $(S_{11} > sthr, \ P_{11} < pthr)$

For each quadrant, a statistic $S_{ij}$ and an error rate $p_{ij}$ is computed. $S_{ij} > 3$ and $p_{ij} < 0.1$ are the thresholds used on the BooleanNet statistics to identify Boolean implication relationships (BIRs). False discovery rate is computed by randomly shuffling each gene and computing the ratio of the number of Boolean implication relationship discovered in the randomized dataset and original dataset. The false discovery rate for BE and EAC dataset was less than 0.001.

Boolean Implication analysis looks for invariant relationship across all the different types of samples regardless of the conditions and treatment protocols. Therefore, it does not distinguish the sample types when discovering Boolean implication relationships. We assume that there are fundamental invariant Boolean implication formula that are satisfied by every sample regardless of their type.

*Construction of BE/EAC Boolean Implication Networks*

A Boolean implication network (BIN) is created by identifying all significant pairwise Boolean implication relationships (BIRs)[4, 10]. The Boolean implication network contains the six possible Boolean relationships between genes in the form of a directed graph with nodes as genes and edges as the Boolean relationship between the genes. The nodes in the BIN are genes and the edges correspond to BIRs. Equivalent and Opposite relationships are denoted by undirected edges and the other four types (low => low; high => low; low => high; high => high) of BIRs are denoted by having a directed edge between them. The network of equivalences seems to follow a scale-free trend; however, other asymmetric relations in the network do not follow scale-free properties. BIR is strong and robust when the sample sizes are usually more than 200 (from our experience of using Boolean Implication for more than 10 years). All our previous papers use thousands of diverse samples to establish Boolean implication relationships. However, Boolean Implication analysis is carried out in such low number of samples such as the selected [GSE100843](GSE100843) (n =76) and [E-MTAB-4054](E-MTAB-4054) (n = 63) datasets. We have demonstrated that we have a reasonable False Discovery Rate ($< 0.001$) when $S > 3$ and $p < 0.1$ are used. Both [GSE100843](GSE100843) and [E-MTAB-4054](E-MTAB-4054) dataset were prepared for Boolean analysis by filtering genes that had a reasonable dynamic range of expression values. When the dynamic range of expression values was small, it was difficult to distinguish if the values were all low or all high or there were some high and some low values. Thus, it was determined to be best to ignore them during Boolean analysis. The filtering step was performed by analyzing the fraction of high and low values identified by the StepMiner algorithm [11]. Any probe set or genes which contained less than 5% of high or low values were dropped from the analysis.

*Generation of Clustered Boolean Implication network*

Clustering was performed in the Boolean implication network to dramatically reduce the complexity of the network. A clustered Boolean implication network (CBIN) was created by clustering nodes in the original BIN by following the equivalent BIRs. One approach is to build connected components in an undirected graph of Boolean equivalences. However, because of noise, the connected components become internally inconsistent e.g., two genes opposite to each other become part of the same connected component. In addition, the size of clusters became unusually big with almost everything in one cluster. To avoid such a situation, we need to break the component by removing the weak links. To identify the weakest links, we first computed a minimum spanning tree for the graph and computed the Jaccard similarity coefficient for every edge in this tree. Ideally if two members are part of the same cluster, they should share as many connections as possible. A threshold is considered for the Jaccard similarity coefficient (0.8 for BE network and 0.5 for the EAC network) below which the edges are dropped from further analysis. Thus, many weak equivalences were dropped using the above algorithm leaving the clusters internally consistent. We removed all edges that have Jaccard similarity coefficient less than the selected threshold and built the connected components with the rest. The connected components were used to cluster the BIN which is converted to the nodes of the CBIN. The choice of the threshold on the Jaccard similarity coefficient play an important role in determining the size and the number of clusters as well as whether they are internally consistent. A new graph was built that connected the individual clusters to each other using Boolean relationships. The link between two clusters (A, B) was established by using the top representative node from A that was connected to most of the members of A and sampling 6 nodes from cluster B and identifying the overwhelming majority of BIRs between the nodes from each cluster.

A CBIN was created using the [GSE100843](GSE100843) and [E-MTAB-4054](E-MTAB-4054) datasets. The edges between the clusters represented the Boolean relationships that are color-coded as follows: orange for low => high, dark blue for low => low, green for high => high, red for high => low, light blue for the equivalent and black for the opposite. A subnetwork is selected using low=>low (blue), high => low (red) and opposite (merged with high=>low as red) edges among the top 10 clusters.

*Charting Boolean paths*

Boolean paths have been explored before to predict the underlying time series events in biological processes such as B cell differentiation [9, 12] and early differentiation events in cancer stem cell [5-7, 13]. This algorithm is called MiDReG (Mining Developmentally Regulated Genes) that uses two seed genes to identify intermediate genes in a biological process.

MiDReG infer intermediate states using a sequence of asymmetric BIRs. Here, using MiDReg algorithm/concept to traverse the Boolean Implication network that identifies paths of clusters where the start and end clusters in the clustered Boolean implication network mark the end points of a possible set of events from healthy to disease. The asymmetric BIRs provide a unique dimension to the network that is fundamentally different from any other gene expression networks in the literature. Traversing a set of nodes in a directed graph of the Boolean network constitutes a Boolean path. A simple Boolean path involves two nodes and the directed edge between them. A complex Boolean path involves more than two nodes and the edges between them.

*Ordering samples based on composite score of Boolean path*

A Boolean path contains one or more clusters. A composite score is computed for each cluster and combined later. To compute the final score, first the genes present in each cluster were normalized and averaged. Gene expression values were normalized according to a modified Z-score approach centered around StepMiner threshold (formula = (expr - SThr)/3/stddev). A weighted linear combination of the averages from the clusters of a Boolean path was used to create a score for each sample. The weights along the path either monotonically increased or decreased to make the sample order consistent with the logical order based on BIR. The samples were ordered based on the final weighted and linearly combined score. A cluster highly expressed in a disease setting received a weight of 1 and healthy setting received a weight of -1.

*Summary of genes in the clusters*

Reactome pathway analysis of each cluster along the top continuum paths was performed to identify the enriched pathways[14]. The pathway description was used to summarize at a high-level what kind of biological processes are enriched in a particular cluster. These can be accessed in **Supplementary Information 2** (for BE) and **Supplementary Information 3** (for EAC).

*Measurement of classification strength or prediction accuracy*

Receiver operating characteristic (ROC) curves were computed by simulating a score based on the ordering of samples that illustrates the diagnostic ability of binary classifier system as its discrimination threshold is varied along with the sample order. The ROC curves were created by plotting the true positive rate (TPR) against the false positive rate (FPR) at various threshold settings. The area under the curve (often referred to as simply the AUC) is equal to the probability that a classifier will rank a randomly chosen IBD samples higher than a randomly chosen healthy samples. In addition to ROC AUC, other classification metrics such as accuracy ((TP + TN)/N; TP: True Positive; TN: True Negative; N: Total Number), precision (TP/(TP+FP); FP: False Positive), recall (TP/(TP+FN); FN: False Negative) and f1 (2 * (precision * recall)/(precision + recall)) scores were computed. Precision score represents how many selected items are relevant and recall score represents how many relevant items are selected. Fisher exact test is used to examine the significance of the association (contingency) between two different classification systems (one of them can be ground truth as a reference).

*AI guided discovery of Boolean paths*

A Boolean path is converted to a path score as mentioned above using a linear combination of normalized gene expression values. The strength of classification of normal and BE/EAC samples using this score is computed by the ROC-AUC measurement. We ranked the clusters based on the ROC-AUC values in the cohort (GSE39491, n = 80, 40 normal, 40 BE) for BE network and E-MTAB-4054 (27 BE vs 17 EAC). Multivariate regression is performed to select the best clusters. The clusters for BE network are filtered by enrichment of differentially expression genes from a recently published BE model (GSE153129). Only two clusters (*SPINK7* and *SLC44A4*) are selected based on this filter: SPINK7 cluster is down-regulated in BE and SLC44A4 cluster is up-regulated in BE.

*Training and Validation Datasets*

A Boolean path is selected after machine learning to construct a Boolean model. The Boolean model is tested in several human and mouse datasets, each comprised of a heterogeneous collection of samples (as mentioned in **Supplementary Information 1**) to demonstrate reproducibility. Selected Boolean path score is computed as mentioned in section "*Ordering* *samples based on composite score of Boolean path*". The sample order using the Boolean path score is evaluated using the sample annotation (normal vs BE; BE vs EAC) by ROC-AUC analysis. We tested how the SPINK7-SLC44A4 path score with weight -1, 1 respectively distinguishes normal and BE samples as they are annotated in training datasets (GSE100843, GSE39491), and validation datasets (GSE65013, GSE64894, GSE49292, GSE26886, GSE34619, GSE13083, E-MTAB-4054). Top two EAC network clusters are selected based on the best ROC-AUC values using the training cohort (E-MTAB-4054): IL10RA, LILRB3. Both these clusters are upregulated in EAC compared to BE samples. The IL10RA-LILRB3 path score is tested with weight 1, 1 respectively to distinguish BE vs EAC samples in training dataset (E-MTAB-4054) and validation datasets (GSE26886, GSE37200, GSE77563). The EAC score is also tested to distinguish normal vs EAC+GEJAC in GSE74553 cohort. We have collected publicly available gene expression datasets derived from mouse models of BE/EAC (GSE24931, GSE103616, GSE158116; **Supplementary Data 1**) to test whether human Boolean models perform well in mice. The gene name conversion from human to the mouse is performed using human genome GRCh38.95 ensembl IDs and mapping data exported from ensemble BioMart web-interface.

*Mathematical model of NE-BE-EAC Progression*

MiDReG[9] (Mining Developmentally Regulated Genes) algorithm is used to model disease progression using concept of invariant Boolean implication formula. A Boolean implication formula is considered invariant if the formula is consistent in almost all samples from a particular domain (almost no exceptions). We observed a Boolean implication formula SPINNK7 high => SLC44A4 low, that is consistent in most diverse global human GSE119087 (n = 25955) dataset. We assumed that this invariant will also hold in NE, BE and EAC samples. When we focus only on the NE, BE and EAC samples, we observed the Boolean implication CXCL8 high => SLC44A4 high (S > 2, p < 0.1; FDR < 0.001) in multiple independent cohorts [E-MTAB-4054, and Pooled GSE26886 (n=58) + GSE40220 (n=3) + GSE42363 (n=14)]. However, combined NE (n=20), BE (n = 20), EAC (n=35), and ESCC (n = 407) samples pooled from GSE26886 (n=67), GSE40220 (n=3), GSE42363 (n=14), GSE69925 (n=266), GSE77861 (n=7), GSE161533 (n=28), GSE32701 (n=29), GSE106185 (n=23), GSE45670 (n=28), GSE17351 (n=5), GSE44021 (n=6), GSE100942 (n=4) and GSE33810 (n=2) failed to show Boolean implication CXCL8 high => SLC44A4 high (S = -0.76, p = 0.91). ESCC samples populate in the CXCL8 high and SLC44A4 low quadrant. All of the pooled samples were microarray datasets generated using the [HG-U133_Plus_2] Affymetrix Human Genome U133 Plus 2.0 Array GPL570 platform. The pooled samples are re-normalized together using RMA algorithm[15]. The NE, BE, and EAC samples nicely organize themselves in three different quadrants (CXCL8 low and SLC44A4 low, CXCL8 low and SLC44A4 high, CXCL8 high and SLC44A4 high) respectively. Matched normal or adjacent non-tumor samples were excluded from the pooled datasets because of contamination from tumor samples (many of them have high levels of CXCL8 expression patterns). Only two normal samples behaved as outliers removed from the analysis (GSM661768 behaved like BE and GSM661777 behaved like ESCC samples). Mathematical model of invariant makes it clear that BE must precede EAC during disease progression.

**Outcome studies**:

*Patient cohort*

We retrospectively analyzed a previously assembled dataset[16] of patients with biopsies reporting BE between 2013 and 2017 and with a complete blood count within 6 months from the endoscopy, as well as patients with esophageal adenocarcinoma (EAC). Cases (n = 113) were classified as non-dysplastic BE (NDBE, n = 72), dysplastic BE (DBE, n = 11) and EAC (n = 30).

Briefly, to enroll the patients with BE, medical records of all patients undergoing upper endoscopy at a tertiary care center in Brazil (Hospital de Clínicas dePorto Alegre) between January 2013 and September 2017 who had columnar epithelium visualized in the distal esophagus were retrospectively analyzed. All endoscopic examinations that fulfilled criteria were included, even though belonging to the same patient in a different period of the surveillance. Exclusion criteria were absence of confirmed intestinal metaplasia on histology (goblet cells on Alcian-Blue staining), immunosuppression by drugs or chronic diseases, active or recent (< 6 months) infectious disease, history of cancer or any hematological or autoimmune disease, and previous surgery of the gastrointestinal tract (except fundoplication for GERD). To be considered eligible, cases also must have had a complete blood count (CBC) collected between the period of 6 months before and 6 months after endoscopy and necessarily outside of a context of clinical emergency (for example, on the emergency room, for any reason) or invasive procedures (such as surgery or esophageal dilation). Those criteria were assessed through a thorough exam of electronic medical records. In the presence of more than one eligible CBC, the one closest to the day of the endoscopy was selected. If an included patient had any endoscopy performed before 2013, those exams were also analyzed for inclusion according to the criteria stated above.

Patients with EAC were selected from hospital discharge diagnostic records between January 2005 and December 2017 if they had the following codes, according to the international classification of diseases (ICD-10; C15: malignant neoplasm of the esophagus): C15.2 (abdominal esophagus), C15.5 (lower third), C15.8 (overlapping sites), and C15.9 (unspecified). Squamous cell carcinoma and gastroesophageal junction (GEJ) tumor types 2 or 3 of Siewert classification14 were excluded. Same exclusion criteria used for BE cases were applied, except the need of confirmed intestinal metaplasia.

Demographic, clinical and endoscopic data collected (as published before[16]). Data collected from endoscopic histopathological reports were histologic diagnosis and adherence to Seattle Protocol, that was evaluated comparing endoscopic descriptions with pathology reports (adherence was considered if sent to pathology separated samples with at least 4 biopsy fragments for every 2 cm of columnar epithelia). Histopathological diagnosis of intestinal metaplasia, dysplasia, and adenocarcinoma was carried out by expert pathologists in the institution in a clinical routine fashion.

All cases were classified into three groups based on histopathological diagnosis: non-dysplastic BE (NDBE), dysplastic BE (DBE)—with either low- or high-grade dysplasia—and esophageal adenocarcinoma (EAC). Staging of EAC patients was done based on TNM classification according to the 7th edition of American Joint Committee on Cancer (2010). EAC patients were divided in 2 groups, according to TNM (stage I/II and III/IV). For patients submitted to esophagectomy without neoadjuvant treatment, pathological stage was used; clinical staging was considered for the rest.

The primary endpoint was the correlation of changing absolute neutrophil count (ANC) with advancing stages of BE progression. Subsequently, ANC was used as a parameter to stratify patients and analyze its impact on clinical outcome. The primary clinical outcome was overall survival and recurrence-free survival (period January 2005 –December 2021).

The study was approved by the Brazilian National Committee on Research Ethics (CONEP), registered by number CAAE-81068617.2.0000.5327.

*Univariate and multivariate analyses*

Prediction of progression from NDBE to DBE to EAC is analyzed using univariate and multivariate analyses based on several clinical parameters such as SEX, ALC, ANC, and PLAT. Univariate and multivariate analyses is performed using Ordinary Least Squares regression (OLS) python statsmodels (version 0.12.2) package.

*Survival analysis in TCGA-EASC and TCGA-STAD dataset*

TCGA-EASC and TCGA-STAD datasets were used to study the relationship between the composite expression score of different genes (or single CXCL8 gene) from various signatures and patient clinical outcomes for esophageal adenocarcinoma (EAC), esophageal squamous cell carcinomas (ESCC) and gastric adenocarcinomas (GC). We evaluated the prognostic value of mRNA expression of either single (CXCL8) or several gene combinations according to overall survival (OS) in cancer patients with high mutation load. Nonsynonymous mutations were discovered using 'missense-variant', 'splice-region-variant', 'splice-donor-variant', 'frameshift-variant', 'stop-gained', 'inframe-deletion', 'splice-acceptor-variant', 'coding-sequence-variant', 'non-coding-transcript-exon-variant', 'non-coding-transcript-variant', 'inframe-insertion', 'start-lost', 'stop-lost', 'NMD-transcript-variant', 'protein-altering-variant' and 'incomplete-terminal-codon-variant' annotations. High mutation load is computed using median threshold of the number of mutations observed in tumor samples based on four different mutation callers from the TCGA portal summarized in UCSC Xena browser: VarScan, MuSE, MuTect, and SomaticSniper. A total of 65 patients with EAC, 26 patients with ESCC and 170 patients with GC were included in all analyses and events were plotted for a 4-year period (1460 days). StepMiner tool was used to compare the predictive value of several gene combinations in patients with low and high expressions. Patients were divided into two groups, high vs. low expression, based on the StepMiner threshold +/- noise margin of gene expression or the composite expression score. Noise margin of a composite signature is estimated by the square root of the sum of squares of the scaled down (1/3/stddev) version of the individual noise margin of the gene which is 0.5 based on the BooleanNet approach. The OS Kaplan-Meier plots are presented with the log-rank *p*-value. For the composite signature scores of CD16 (FCGR3A and FCGR3B) in EAC, threshold was computed by using StepMiner twice, first on the whole scores and then using all the values lower than the first StepMiner threshold. For ESCC, threshold for composite score of CD16 (FCGR3A and FCGR3B) was computed by using StepMiner twice, first on the whole scores and then using all the values greater than the first StepMiner threshold.

### **Genomic analyses**:

*Patient samples*

Whole-genome sequencing data for 320 esophageal biopsies from 80 patients with Barrett's esophagus (BE) were analyzed (de-identified data publicly available, dbGaP Study Accession**:** phs001912 and phs001654). Longitudinal data from patients were collected as part of a case-control study design performed at the Fred Hutchinson Cancer Research Center. Demographics for patient cohort have been previously described[17]. Progressors (cases) were defined as patients who progressed to esophageal adenocarcinoma (EAC) during surveillance (n=40 total) in the Seattle Barrett's Esophagus Program. Non-progressors (controls) were defined as those who did not progress to EAC throughout long-term follow-up (n=40 total).

*Mutational analysis*

After downloading the BE genomics data, quality control checks were performed for each sample to remove any sequencing files that do not meet our stringent criteria. Briefly, samples were greylisted when: (i) FASTQ file(s) failed one or more of the criteria from FastQC1; (ii) a tumor-normal pair had less than 95% concordance as estimated by Conpair2; (iii) samples exhibited low tumor purity, as estimated by ascatNgs3; (iv) samples exhibited low sequencing coverage for either the cancer or the normal tissue. Additionally, after mutation calling was completed, any sample with more than 10% of mutations attributed to signatures of known sequencing artifacts was greylisted. Only two of the original BE samples failed these quality control checks. Note that while somatic mutations and germline variants were identified in all greylisted samples, these samples were not used for any subsequent analyses. Somatic mutations and germline variants were identified using our ensemble variant calling pipeline. Specifically, FASTQ files were aligned to human genome build GRCh38.d1.vd1 using BWA-MEM5. Duplicate reads were marked using Picard MarkDuplicate6. The alignment was further refined through local indel realignment and base score recalibration using the GATK toolkit7,8. Next, the paired cancer and normal BAM files were subjected to somatic mutation calling analysis using four state-of-the-art computational tools. More specifically, single base substitutions (SBSs) and small insertions and deletions (indels) were identified independently using Mutect29, VarScan210, Strelka211, and MuSE12. Germline single-nucleotide polymorphisms (SNPs) were detected using VarScan2. Copy-number alterations were identified using ascatNgs3. Somatic mutations marked as "PASS" by Mutect2 and Strelka2 were filtered based on their mutation confidence scores: TLOD score >= 10 (Mutect2) and SomaticEVS >= 15 (Strelka2). Any somatic mutations found in 2 or more of the 4 variant callers were considered as high confidence mutations; this optimum combination strategy is based on the previous experience from pan-cancer analysis performed by TCGA MC313 and ICGC PCAWG14. Subsequent filters were used to produce the final set of mutations for downstream analysis. The following filters addressed most artifact mutations introduced during sample preparation and/or sequencing: (i) DKFZbiasFiter15 removed SBSs with biased variant read support, which can be a result of unbalanced PCR amplification, biased reading during sequencing, and DNA shearing leading to 8-oxoguanine lesions during sample preparation16; (ii) panel of normal (PON) filter, generated using Mutect2, removed additional technical artifacts recurrently appearing in genomic locations of normal samples; (iii) variant allele frequency threshold was applied to all SBSs and indels to remove any low frequency mutations, which were commonly introduced as part of the DNA sequencing process17. Overall, this analysis generated a catalog of harmonized bona fide DNA somatic mutations and germline variants in each BE sample.

For downstream analyses, we further filtered for somatic mutations from this catalog that were found by all 4 callers described above. For each patient, we then considered the set of unique somatic mutations across their 4 samples from 2 timepoints (including 2 sampling locations of the upper esophagus and lower esophagus regions at each timepoint).

*Statistical analysis*

To calculate the number of occurrences of mutations in the cluster of genes identified from the BE→EAC map (**Figure S7A**), we cycled through merged VCF files from EAC Progressors (n=40) and summed the total number of mutations in each patient across the genes in each cluster as shown in blue boxplots. The same calculations were performed for non-progressor patients (n=40) and total mutations are shown in red boxplots. P-values from two-sided Mann–Whitney *U*-tests are reported above each plot comparing Progressor and Non-progressor patients for each cluster (statistical tests performed using scipy.stats.mannwhitneyu in Python v3.9.7).

To calculate the number of occurrences of mutations in neutrophil function associated genes, we cycled through the merged VCF files from patients who were EAC Progressors (n=40) and summed the total count for each gene. These totals by gene were then compared to totals counted in the same way within the non-progressor patients (n=40), see **Figure S7B**.

### **Experimental Approaches:**

*Reagents and antibodies*

Antibodies that were used in this work include rabbit serum anti-SUPT6H (ThermoFisher, #A300-801A), and rabbit serum anti-P63/TP63 (BosterBio, #PA2056). ImmPRESS HRP Horse Anti-Rabbit IgG Polymer Detection Kit Peroxidase was purchased through Vector Laboratories (#MP-7401-50), and the 2-Component DAB pack was purchased from BioGenex (#HK542-XAKE), as was the hematoxylin (#HK100-9K).

*Human subjects with or without BE*

Esophageal biopsies used for IHC were obtained from patients undergoing endoscopies as a part of their routine care and follow-up at UC San Diego's Center for Esophageal Diseases. Patients were recruited and consented using a study proposal approved by the Institutional Review Board of the University of California, San Diego Center for Esophageal Diseases GEODE (Gastro-Esophageal Oncogenesis, Dysmotility, & Evolution) Research Program, Division of Gastroenterology, University of California San Diego, following the protocol approved by the Human Research Protection Program (HRPP) Institutional Review Board (Project ID#200047). Mucosal biopsies were obtained during sedated upper gastrointestinal endoscopy using cold forceps from two cohorts: 1) Asymptomatic healthy, 2) BE. For all cohorts, biopsies were obtained from the squamous epithelium 5 cm proximal to the Z line. Additionally, for the BE cohort the BE segment was closely inspected under high-definition white light and narrow band imaging, and biopsies were obtained from the esophagus 2-5 cm proximal to the BE segment. The clinical phenotype and information were curated based on histopathology reports from Clinical Pathology and Chart check. For all the deidentified human subjects the information including age, ethnicity, gender, previous history of disease and medication were collected from the chart following the security and privacy rules outlined in the HIPAA (Health Insurance Portability and Accountability Act of 1996) legislation. Written informed consent was obtained from all participants. The study design and the use of human study participants was conducted in accordance with the criteria set by the Declaration of Helsinki.

*Immunohistochemistry of patient esophageal samples*

Formalin-fixed, paraffin-embedded (FFPE) tissue sections of 4 µm thickness were cut and placed on glass slides coated with poly-L-lysine, followed by deparaffinization and hydration. Heat-induced epitope retrieval was performed using tris EDTA (pH 9.0) in a pressure cooker using a rolling boil for 15 minutes. Tissue sections were incubated with 3% hydrogen peroxidase for 5 minutes to block endogenous peroxidase activity, followed by a one-hour incubation with 2.5% horse serum. Slides were then incubated with primary antibodies for 1.5 hours in a humidified chamber at room temperature. After primary incubation, slides were incubated with secondary antibodies (horse, anti-rabbit) for 30 minutes at room temperature and washed. Antibodies used for immunostaining; SUPT6H [1:250], TP63 [1:500]. Immunostaining was visualized using 3,3′-diaminobenzidine chromogen and counterstained with Mayer's Hematoxylin. Samples were quantitatively analyzed and scored based on the presence (positive) or absence (negative) of staining.

*IHC Quantification*

IHC images were randomly sampled at different 300x300 pixel regions of interest (ROI). The ROIs were analyzed using IHC Profiler[18]. IHC Profiler uses a spectral deconvolution method of DAB/hematoxylin color spectra by using optimized optical density vectors of the color deconvolution plugin for proper separation of the DAB color spectra. The histogram of the DAB intensity was divided into 4 zones: high positive (0 to 60), positive (61 to 120), low positive (121 to 180) and negative (181 to 235). High positive, positive, and low positive percentages were combined to compute the final percentage positive for each region of interest (ROI). The range of values for the percent positive is compared among different experimental groups. Data is displayed as percent positive stain.

*Statistical analysis*

Statistical significance between experimental groups was determined using two-tailed Mann-Whitney test (IHC score). For all tests, a p-value of 0.05 was used as the cutoff to determine significance. All experiments were repeated a least three times, and p-values are indicated in each figure. All statistical analysis was performed using GraphPad prism 8.4.3.

**Supplementary Text**

Not applicable.

**A**

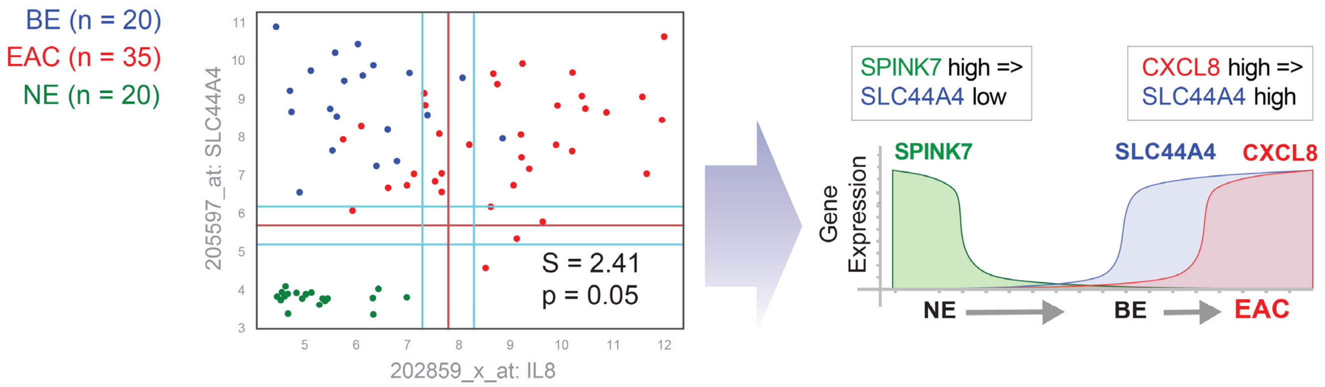


GSE26886 (n=58),
GSE42363 (n=14), GSE40220 (n=3)

**B**

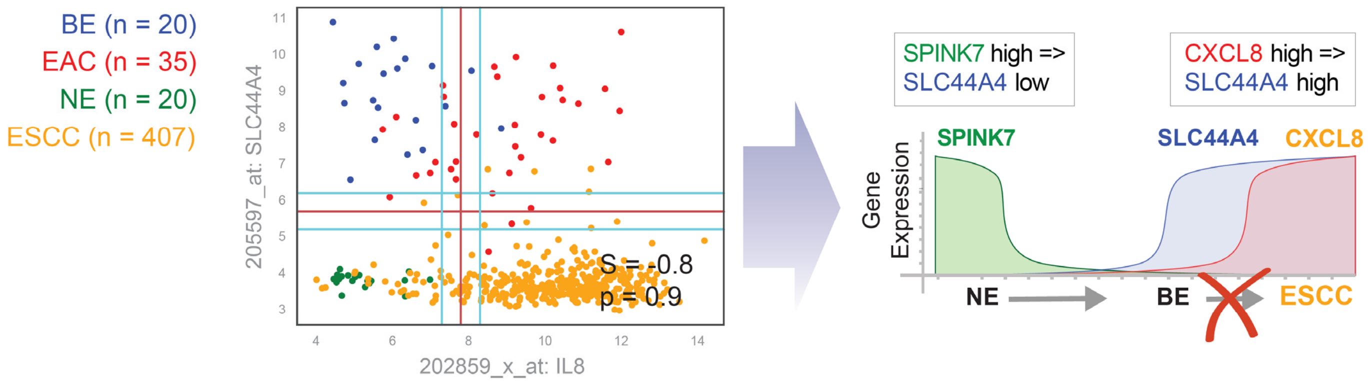


GSE26886 (n=67), GSE40220 (n=3), GSE42363 (n=14),
GSE69925 (n=266), GSE77861 (n=7), GSE161533 (n=28),
GSE32701 (n=29), GSE106185 (n=23), GSE45670 (n=28),
GSE17351 (n=5), GSE44021 (n=6), GSE100942 (n=4),
GSE33810 (n=2)

**Figure S1. A Boolean logical model of NE→BE→EAC progression shows that EACs, but not ESCCs arise from BE. [Related to Figure 5]**

**A.** A scatterplot (left) for *CXCL8* and *SLC44A4* expression in a pooled cohort of dataset (individual GSEID# are indicated beneath the graph). All of them were microarray datasets generated using the [HG-U133_Plus_2] Affymetrix Human Genome U133 Plus 2.0 Array GPL570 platform). *CXCL8* high => *SLC44A4* high (S = 2.41, p = 0.05, FDR < 0.001) in an invariant Boolean implication relationship in NE, BE and EAC samples where each sample type is mostly confined to one quadrant (NE, bottom-left; BE, top-left; and EAC, top-right). Schematic (right) to visualize the mathematical model of NE→BE→EAC progression based on MiDReG[9, 12] analysis using Boolean Implication relationships. Since *CXCL8* high => *SLC44A4* high, and both *CXCL8* and *SLC44A4* are low in NE, the invariant model suggests that BE stage (*SLC44A4* high, *CXCL8* low) must precede *CXCL8* high and *SLC44A4* high.

**B**. A scatterplot (left) generated as in A in a pooled cohort of dataset to include ESCCs (individual GSEID# are indicated beneath the graph). Unlike EACs (A), ESCCs fail to remain confined to the upper right quadrant. All of them were microarray datasets generated using the [HG-U133_Plus_2] Affymetrix Human Genome U133 Plus 2.0 Array GPL570 platform). Schematic (right) to visualize the conclusion of the scatterplot, showing that NE→BE→EAC progression model fails in the case of BE→ESCC progression.

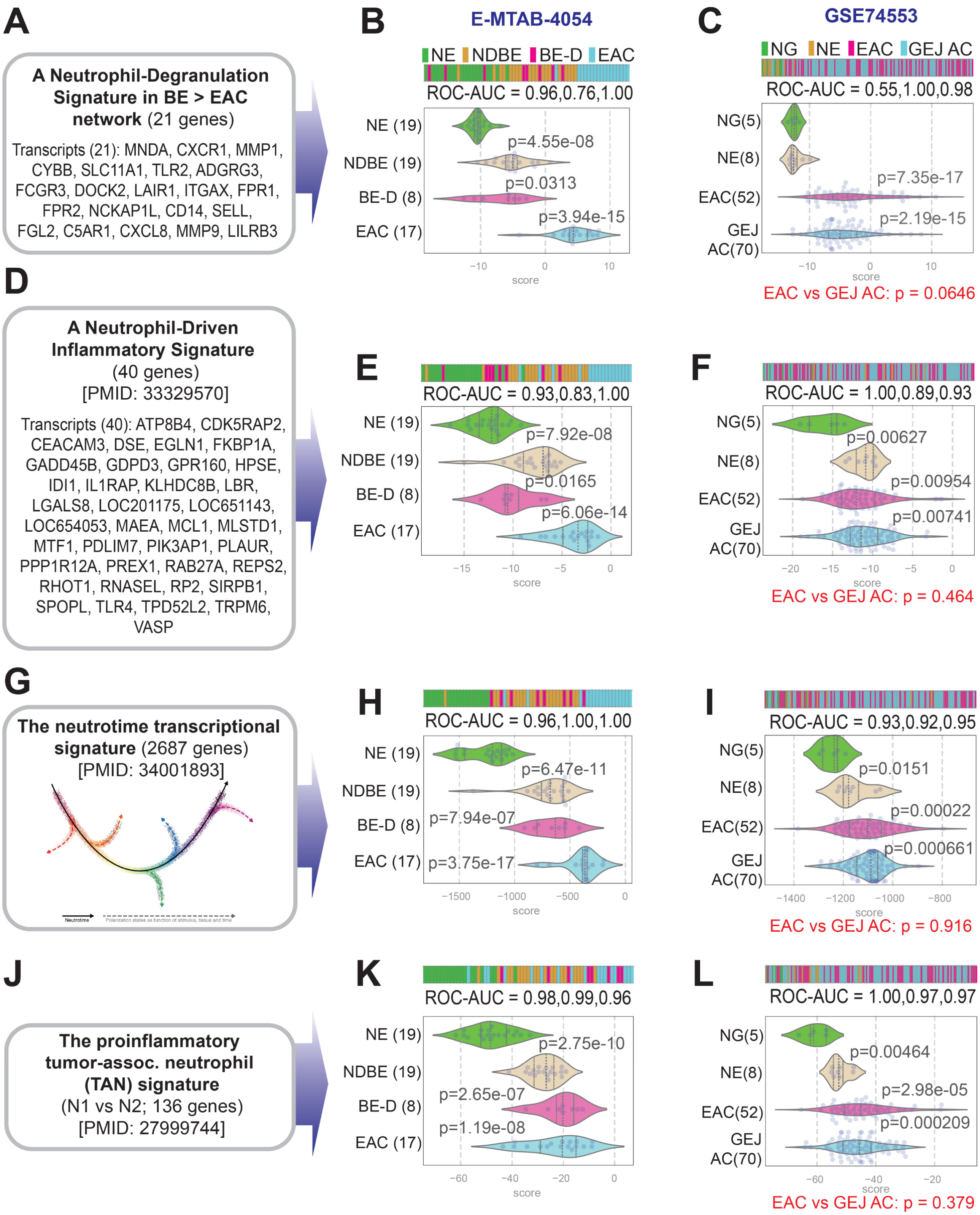


**Figure S2. The EAC and GEJ-AC tumor immune microenvironments are similar, and characterized by neutrophil processes [Related to Figure 5]**

Bar plots (top) and violin plots (bottom) show the composite scores of upregulated gene clusters (panels A, D, J, E) in normal esophagus (NE), non-dysplastic BE (NDBE), dysplastic BE (DBE), and EACs in E-MTAB-4045 (B, E, H, K) and normal gastric (NG), normal esophagus (NE), EAC and GEJ-AC in GSE74553 (C, F, I, L). *P* values indicate comparison of each sample type against the NE (middle) or NG (right) samples, as determined by Welch's t-test. The neutrophil degranulation (ND) signature (A) was derived from the EAC signatures identified in this work.

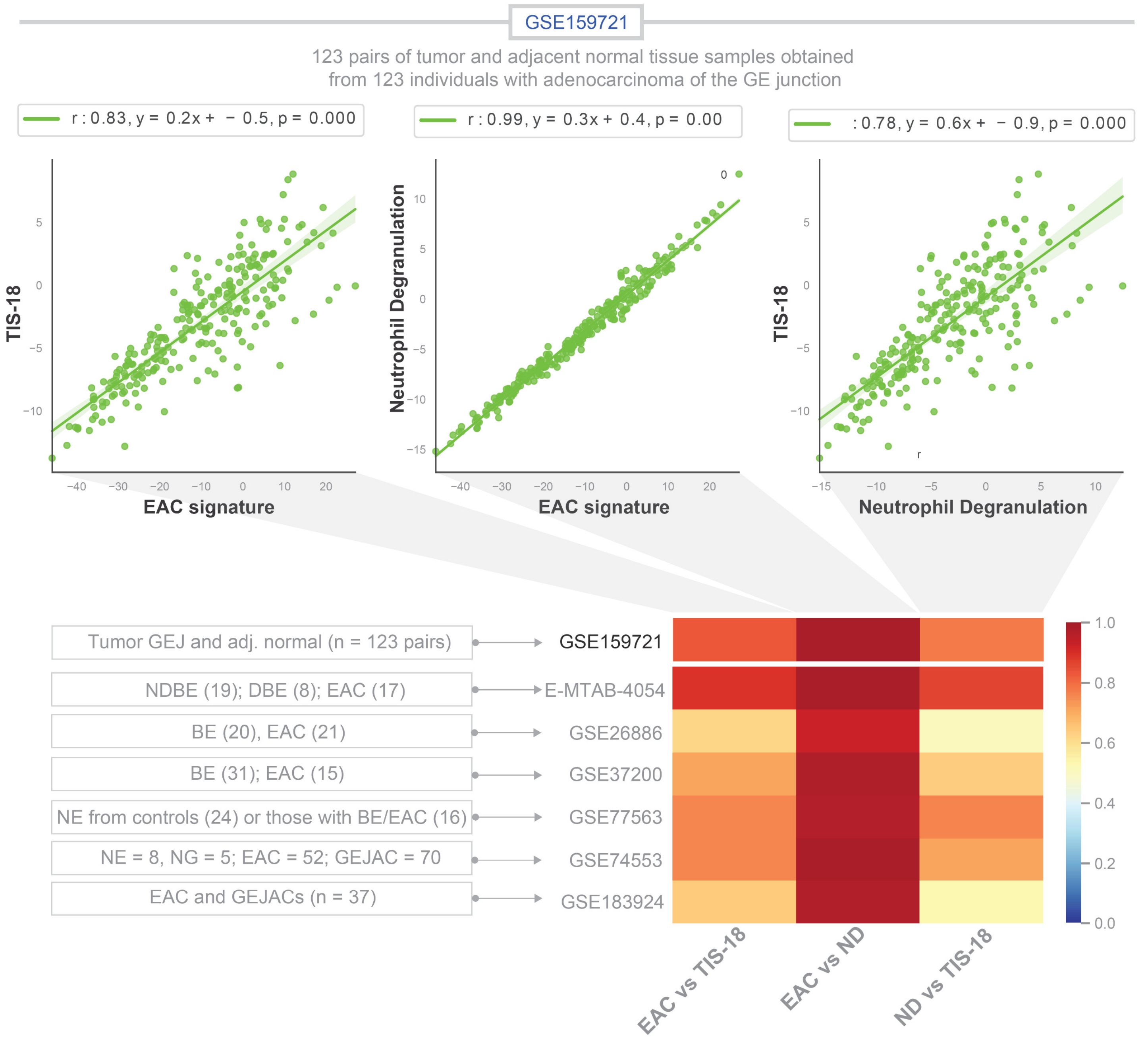


**Figure S3. The neutrophil degranulation signature in EACs and GEJ-ACs correlate positively with tumor inflammatory signature (TIS) which is indicative of benefit from immune checkpoint therapy. [Related to Figure 5]**

Top: Correlation tests were calculated in GSE159721 (123 pairs of normal GEJ and GEJ-ACs) and displayed as scatter plots using python seaborn lmplots with the p-values between the 18-gene TIS signature on the Y axis and EAC signature on the X axis (*left*), neutrophil degranulation signature on the Y axis and the EAC signature on the X axis (*middle*) and the 18-gene TIS signature on the Y axis and neutrophil degranulation signature in the X axis (*right*).

Bottom: “r” values from correlation tests calculated on numerous independent cohorts are displayed as heatmap.

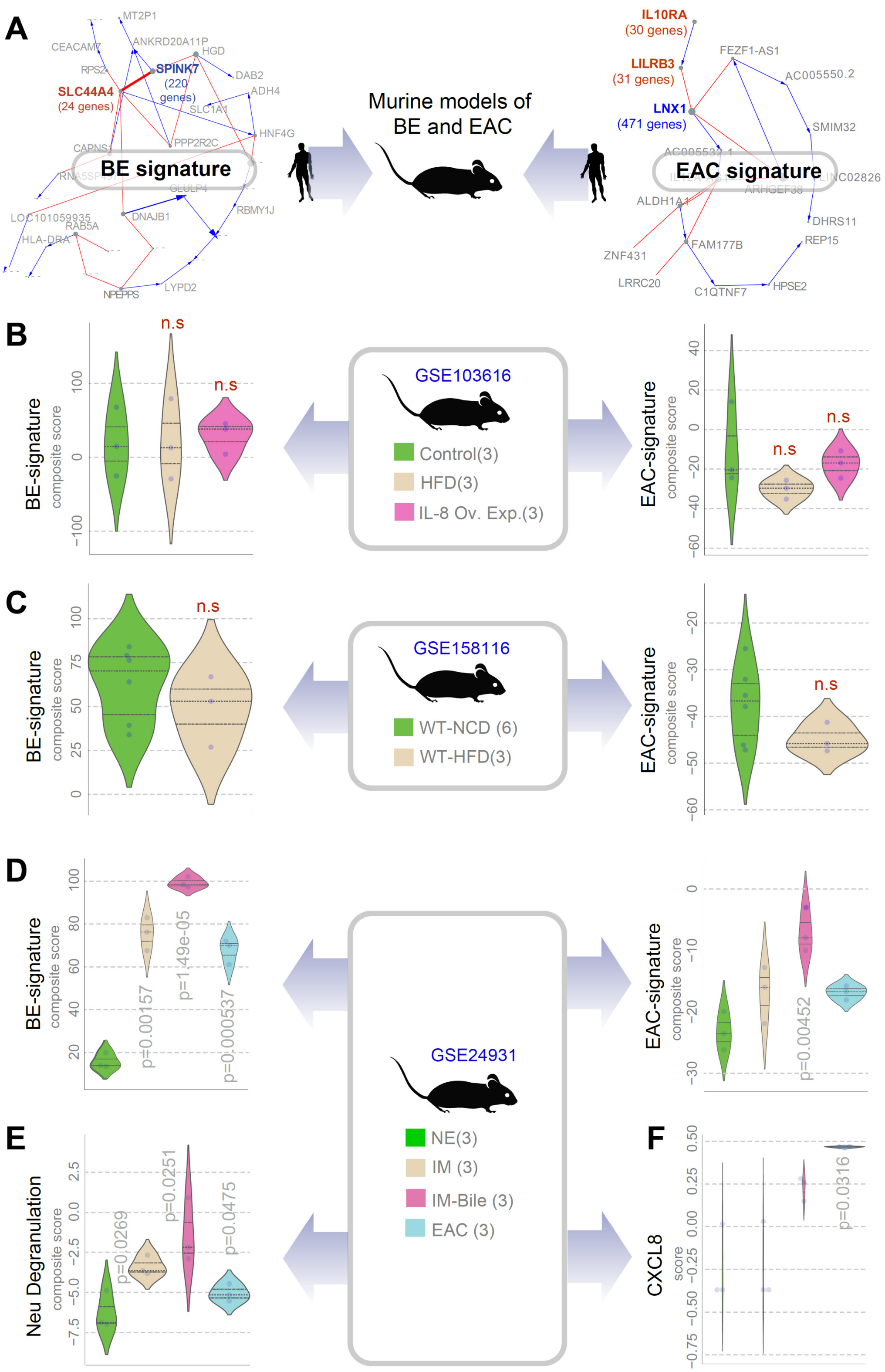


**Figure S4. The tumor immune microenvironment in human EAC is rarely recapitulated in murine EAC models [Related to Figure 5]**

**A.** Schematic displays the general strategy used here to vet the appropriateness of murine models for modeling human diseases using human Boolean map-derived BE/EAC signatures. Gene signatures from these human maps are assessed for induction in the murine datasets.

**B-F**. Violin plots showing the composite scores of upregulated gene clusters (BE, EAC, Neu degranulation) or single gene (*CXCL8*) analyzed in murine models of BE→EAC progression, which includes high fat diet (HFD; B, GSE103616; C, GSE158116) and IL8-overexpression (Ov. Exp.; B, GSE103616. *P* values assessed using Welch's test were all insignificant in B-C, and only significant values are displayed. NE, normal esophagus; IM = intestinal metaplasia.

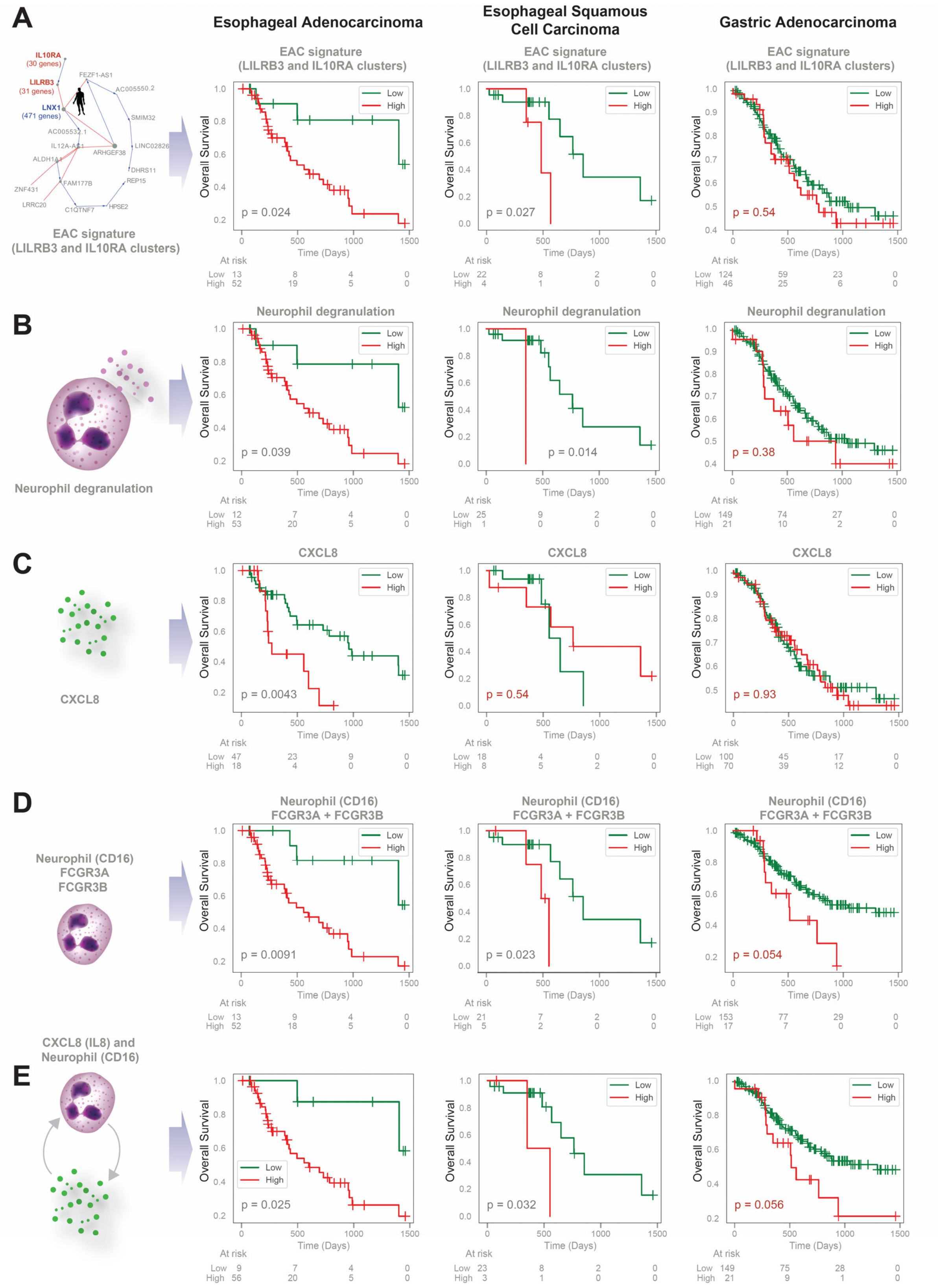


**Figure S5. Prognostic value of gene signatures in EACs, ESCCs and gastric adenocarcinomas (GCs). [Related to Figure 6]**

Kaplan-Meier (KM) plots display the overall survival of patients with tumors stratified based on the high vs low composite scores of gene signatures displayed as schematics on the left. P values were determined by logrank analysis. EAC and ESCC KM plots in panels C and D are repeated from **Figure 6K-L**.

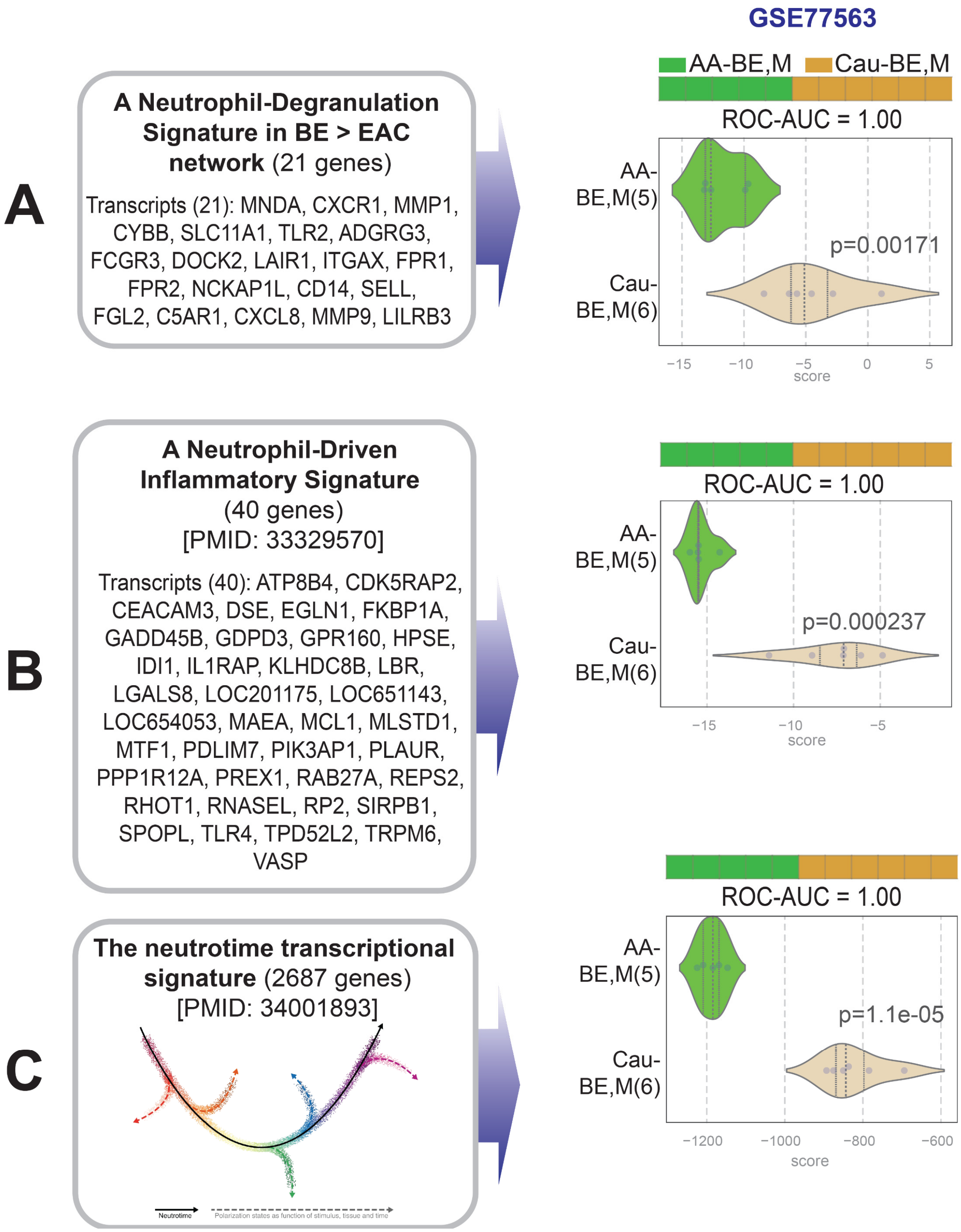


**Figure S6. Neutrophil signatures are differentially induced in the histologically normal esophageal lining proximal to BE/EAC lesions in AA vs Cau subjects.** [Related to Figure 7]

Bar plots (top) and violin plots (bottom) show the composite scores of upregulated gene clusters (panels A-C) in biopsies from histologically normal esophagus from AA vs Cau subjects with the diagnosis of BE/EAC (in GSE77563). Statistical significance was determined by Welch's t-test. The neutrophil degranulation (ND) signature (A) was derived from the EAC signatures identified in this work.

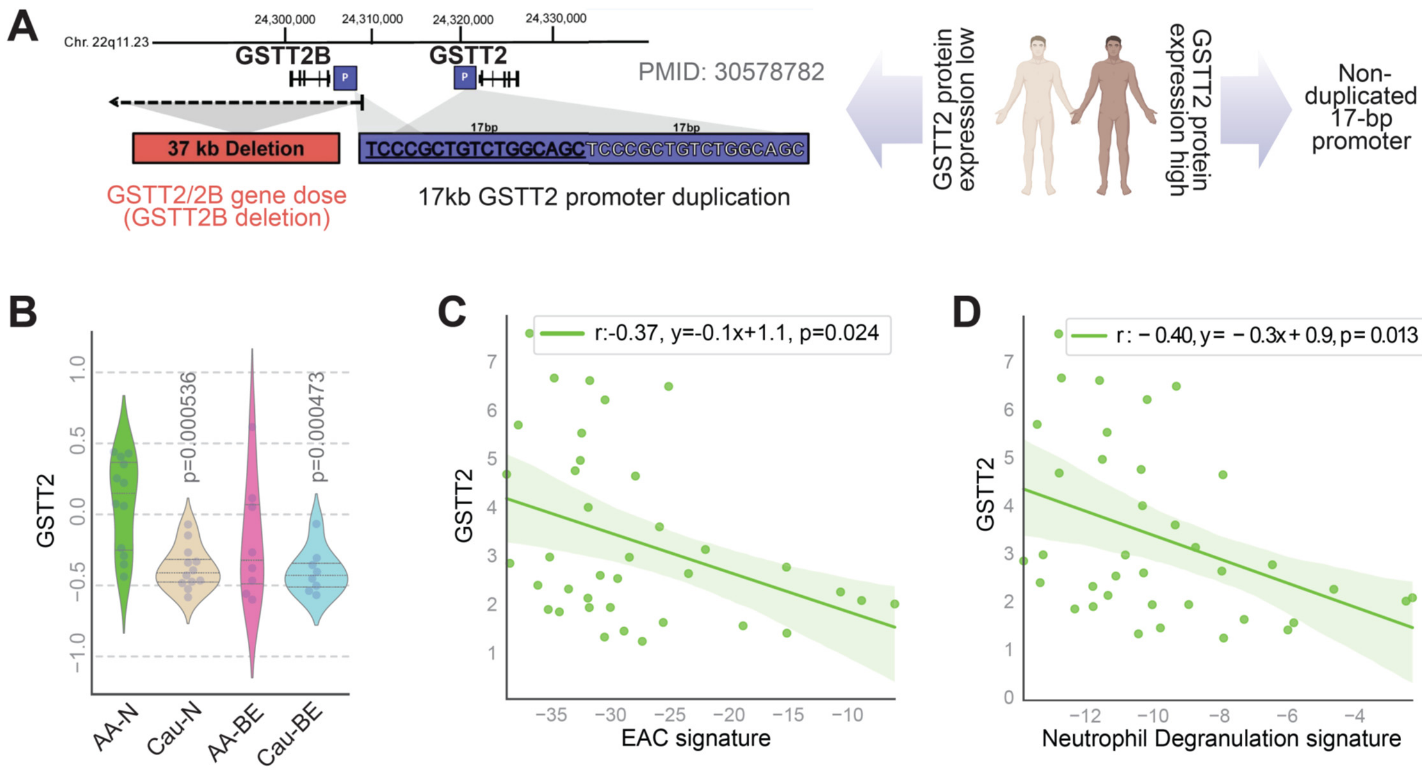


**Figure S7**. **GSTT2 expression inversely correlates with EAC and neutrophil degranulation signatures in the histologically normal esophageal lining proximal to BE/EAC lesions in AA vs Cau subjects.** **[Related to Figure 7]**

**A**. Schematic summarizing the key conclusions drawn in the only other racially influenced determinant of EAC risk in AA vs Cau. Expression of GSTT2, which protects esophageal squamous cells against DNA damage from genotoxic stress is reduced in Cau compared to AA (right) due to genomic variants at the GSTT2 locus, i.e., either a 37-kb deletion and a 17-bp promoter duplication (left).

**B.** Violin plots showing the abundance of GTTT2 in normal esophagus from control (AA-N and Cau-N) BE/EACs (AA-BE and Cau-BE) subjects. *P* values indicate comparison of each sample type against the normal samples, as determined by Welch's t-test.

**C-D**. Correlation tests between GSTT2 on the Y axis and EAC (G) or neutrophil degranulation (H) gene signature scores on the X axis were calculated and displayed as scatter plots using python seaborn lmplots with the p-values. The confidence interval around the regression line is indicated with shades.

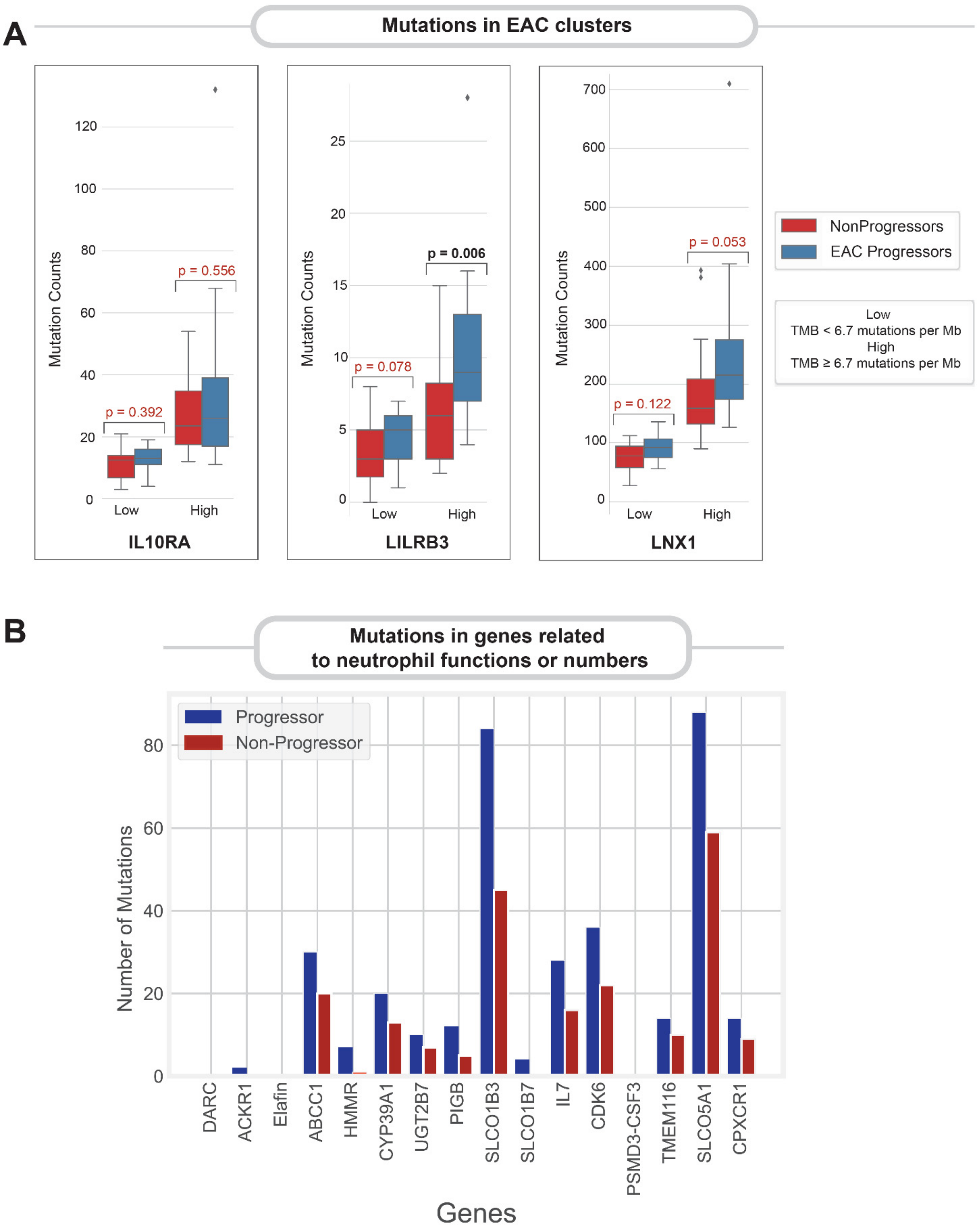


**Figure S8**. **Somatic mutations in genes within the EAC signature or those related to neutrophil function.**

**A. Mutational counts in invariant gene clusters.** Numbers of somatic mutations detected in the 3 gene clusters (identified in the EAC map) in BE patients who progressed to EAC ('EAC Progressors', n=40) and age-matched BE patients who did not ('Non-Progressors', n=40) [data from[17]]. P-values from two-sided Mann–Whitney *U*-tests are reported above each plot comparing Progressor and Non-progressor patients for each cluster.

**B. Somatic mutations in neutrophil genes.** Total mutation counts in neutrophil function associated genes across all Progressor patients (blue bars) vs all non-progressor patients (red bars) [data from[17]]. Consensus mutations required to be called by 4 variant callers (MuTect2, VarScan, Strelka, MuSe).
